# Distinct selection signatures during domestication and improvement in crops: a tale of two genes in mungbean

**DOI:** 10.1101/2022.09.08.506689

**Authors:** Ya-Ping Lin, Hung-Wei Chen, Pei-Min Yeh, Shashi S. Anand, Jiunn Lin, Juan Li, Thomas Noble, Ramakrishnan Nair, Roland Schafleitner, Maria Samsonova, Eric Bishop-von-Wettberg, Sergey Nuzhdin, Chau-Ti Ting, Robert J. Lawn, Cheng-Ruei Lee

## Abstract

Domestication and improvement are two crucial processes underlying the evolution of crops. Domestication transformed wild plants into a utilizable form for humans; improvement refined cultivars adapting to distinct environments and local preferences. Using whole-genome re-sequencing of *Vigna radiata*, we investigated the demographic history and compared the genetic footprints of domestication and improvement. The Asian wild population migrated to Australia at about 50 kya, and domestication happened in Asia about 9 kya selecting for non-shattering pods. The key candidate gene for this trait, *VrMYB26a*, has lower expression in cultivars, consistent with the reduced polymorphism in the promoter region reflecting hard selective sweep. The determinate stems were later selected as an improvement phenotype and associated with the gene *VrDet1*. Two ancient haplotypes reducing gene expression exhibit intermediate frequencies in cultivars, consistent with selection favoring independent haplotypes in soft selective sweep. Our results suggest domestication and improvement may leave different genomic signatures of selection, reflecting the fundamental differences in the two processes and highlighting the limitations of genome-scan methods relying on hard selective sweep.

## Introduction

Crop evolution encompasses two phases, domestication and improvement (Hufford, et al. 2012; Li, et al. 2013; Meyer and Purugganan 2013; Abbo, et al. 2014; Giovannoni 2018; Kumar, et al. 2021). The former refers to the initiation of the divergence from the wild progenitor. Strong selection occurs upon a small population of wild progenitors, with accompanying loss of genetic and phenotypic diversity. Given that domestication aims to increase human benefits rather than plant fitness in the wild, this process typically leads to the so-called domestication syndrome, including the loss of seed shattering and dormancy, gain of gigantism, and the reduction of lateral branches. For example, teosinte (*Zea mays* ssp. *parvigalumis*), the progenitor of maize, has more lateral branches and a tendency for ear shattering than maize (Doebley, et al. 1995). *Oryza rufipogon*, the progenitor of rice, shows more easily seed shattering and smaller seeds than rice (Huang, Kurata, et al. 2012). Improvement, on the other hand, generally occurred accompanying the spread of the domesticated populations to different agro-ecological regions, primarily focusing on increasing yield and the enhanced adaptation to local environments or cultivation systems. For example, soybean varieties in northern Japan required shorter days to flowering than those in southern Japan, suggesting that the regional farmers selected soybeans according to local environments (Kaga, et al. 2012). Given the different natures of these two selection processes, it is intriguing whether they affected the underlying genes and crop genomes in different ways.

Meyer et al. (2013) proposed criteria for defining domestication and improvement genes in crops. Genes controlling domestication traits are supposed to undergo positive selection, and the causal mutations would eventually become nearly fixed in all lineages from a single domestication event (Meyer and Purugganan 2013). For example, *sh4*, the loss-of-function allele of one of the influential domestication genes controlling non-shattering in rice, is almost fixed in cultivated rice (Li, et al. 2006). On the other hand, improvement genes may exhibit different signatures of selection since the target phenotypes for improvement often differ depending on local environments and cultivation systems, diversifying the phenotypes of worldwide varieties (Huang, Zhao, et al. 2012; Zhang, et al. 2015). For instance, in soybean, the mutant allele of the flowering-time gene *E1* resulted in the change of photoperiod requirement and was enriched in specific populations, associated with the adaptation of soybean cultivars to diverse environments (Xia, et al. 2012; Zhou, et al. 2015). Despite the general knowledge of expected differences between domestication and improvement processes, only few have specifically contrasted their genomic signatures of selection in the same species.

*Vigna radiata*, mungbean, a traditional legume crop in Asia, was believed to be originated in India (Purugganan and Fuller 2009). Its putative ancestor, *V. radiata* var. *sublobata*, occupies a wide range from Africa, South Asia, Indonesia, to Australia (Tateishi 1996; Castillo and Fuller 2010). Two recent studies used reduced-representation sequencing to investigate the genetic structure of cultivated mungbean (Noble, et al. 2018; Breria, et al. 2020), but genome-wide investigation comparing the cultivars and wild progenitors is lacking. The genetic architecture of trait differences between wild and cultivar groups were investigated in bi-parental crosses (Isemura, et al. 2012; Li, et al. 2018). Some of these traits are classic domesticated traits, such as the loss of pod twisting and seed dormancy, and other traits, such as plant stature and yield components, resulted from the improvement phase (Huyghe 1998; Fuller and Allaby 2018). To investigate the genomic and genetic patterns of domestication and improvement, studies using whole-genome sequencing from many wild and cultivar accessions are required.

In this study, we re-sequenced 114 *V. radiata* accessions and investigated demographic histories and origins of distinct wild and cultivar groups. We targeted two genes controlling important domestication and improvement traits, pod twisting and stem determinacy, to elucidate how artificial selections during these two processes shape genetic architecture and leave distinct selection signatures at targeted loci.

## Results

### Genetic variation of global *V. radiata*

The whole-genome sequencing data were generated for 114 *V. radiata* accessions (36 wild and 78 cultivars) and one accession of closely-related species, *V. mungo*. The sequencing depth was 33.52 ± 9.76X per accession. The mapping rate of these raw reads to the *V. radiata* reference genome version 6 was 98.08 ± 2.67% (Supplementary Table S1). Finally, we identified 18,548,167 (18.5 million) bi-allelic SNPs for the following analyses.

We used these SNPs to infer the genetic structure. Lower cross-validation error values of ADMIXTURE exist in K = 2 and 5 (Supplementary Fig. S1). Accessions from the same continent were clustered together when K = 2 (Figure 1A separating Asia and Australia; Supplementary Table S2). Under K = 5, the wild accessions were divided into eastern Australian ones (SubAUe), western Australian ones (SubAUw), and Asian ones (SubAS), and the cultivars were separated into two groups from South Asia (RadSA) and other regions in Asia (RadOther). A similar pattern was observed in the phylogenetic tree and network (Figure 1A-B and Supplementary Fig. S2). The mean *F_ST_* value between wild and cultivar populations was 0.49, showing a high degree of differentiation. The wild groups have higher nucleotide diversity than the cultivar groups (Supplementary Fig. S3A). Likewise, LD decays much faster in the wild than in the cultivar population, reaching *r^2^* = 0.2 at about 22 kb and 173 kb, respectively (Supplementary Fig. S3B).

**Figure 1.**
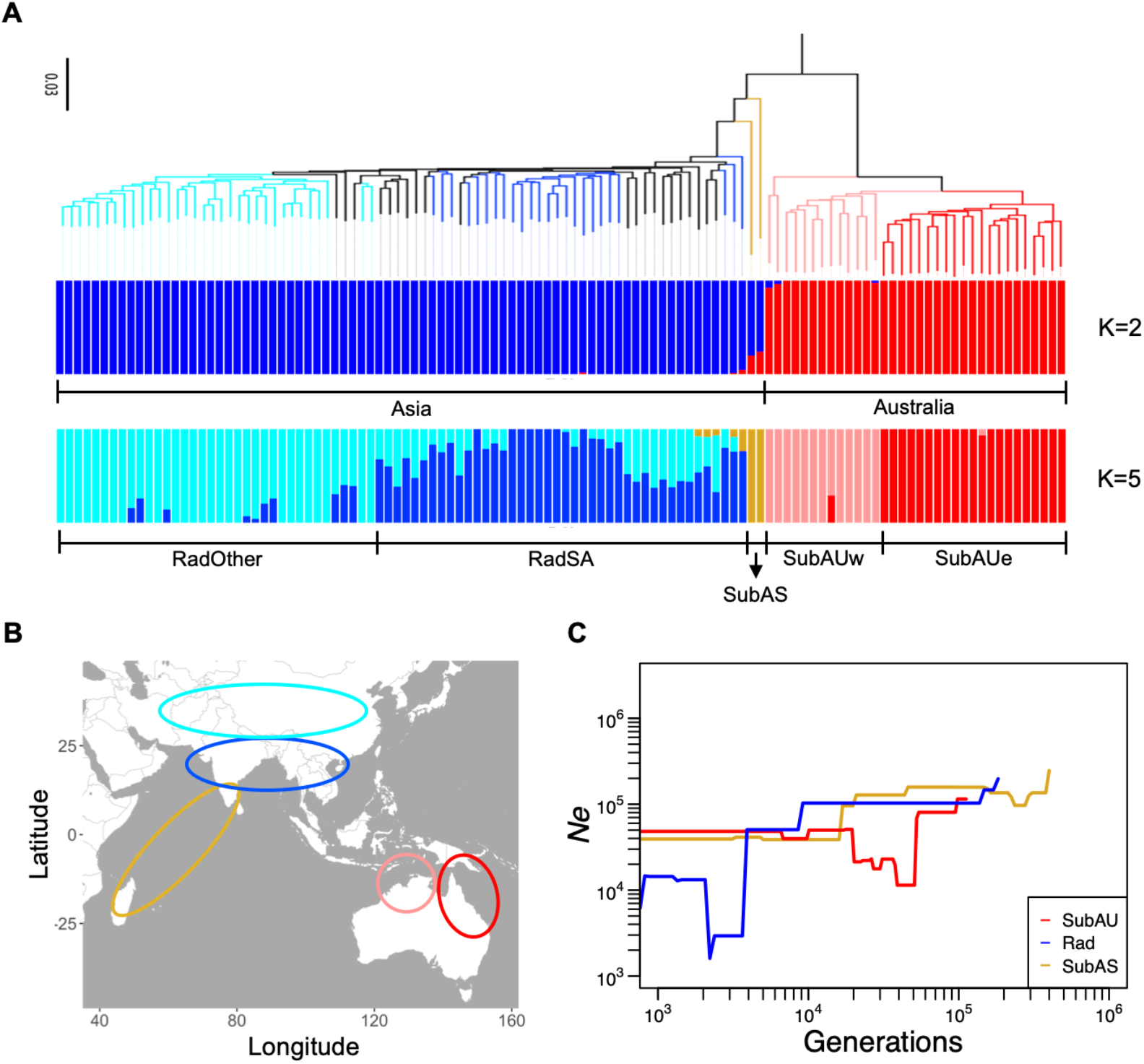
Population structure and demographic history. (A) Neighbor-joining phylogenetic tree and population structure of the 114 mungbean accessions. *Vigna mungo*, VI035226, is the outgroup of the phylogenetic tree. (B) Geographic distribution of the inferred genetic groups. (red: SubAUe, pink: SubAUw, goldenrod: SubAS, blue: RadSA, skyblue: RadOther) (C) Inferred demographic history.

### Demographic history

Most species closely related to *V. radiata* exist in Asia (Norihiko, et al. 2010), so mungbean likely has an Asian origin. The existence of many wild accessions in Australia suggests ancient migration from Asia across the Wallace line. We estimated that the Asian and Australian groups first diverged at about 50 thousand years ago (kya) (Figure 1C, Supplementary Fig. S4A-B). After the divergence, the effective population size of SubAU decreased by about tenfold while that of SubAS remained relatively stable, consistent with SubAU being a founder population colonizing Australia. After this major vicariance between Asia and Australia, Asian cultivars (Rad) diverged from SubAS at about 9 kya (Figure 1C, Supplementary Fig. S4C). After the initial domestication, the effective population size of Rad dropped close to 20 times since 4 kya, presumably due to intensified artificial selection and coinciding with archeological records during 4 to 3 kya (Fuller 2007). The decreased effective population size of Rad might also be affected by climate change, as 4 kya coincides with the start of climate cooling after the Holocene climate optimum (Marcott, et al. 2013).

The strong divergence between Asian and Australian groups was also apparent from the 2D site frequency spectrum comparing Rad, SubAUe, and SubAUw (Supplementary Fig. S5). SubAUe and SubAUw had more shared four-fold degenerate SNPs than each of them with Rad, consistent with their closer relationship. On the contrary, SNPs with high impact were enriched in low-frequency derived-allele categories with few shared between populations. These SNP sites were probably under negative selection, and presumably, detrimental ancestral variations were unlikely to be retained in both descendant populations.

Our results suggested Austalian wild (SubAU) population diverged from the Asian wild group (50 kya) (Figure 1C) during the last ice age (115-11.7 kya), which allows us to track the change in the environmental niche space of these groups. During the last interglacial period (120-140 kya), when *V. radiata* had not colonized Australia, Australia was a less suitable habitat for the Asian groups (Figure 2). During the last ice age, the Sunda Shelf and northern Sahul Shelf were suitable for both the Southeast Asia and Southern Hemisphere populations, facilitating the expansion of *V. radiata* from Southeast Asia into Australia both geographically and environmentally (Figure 2, Supplementary Fig. S6). The Schoner’s D values (higher values representing higher niche similarity) also supported that the Southern Hemisphere population shared more suitable habits with the Southeast Asian population (0.612) than the South Asian one (0.418). At present, most of the Malay Archipelago and Australia appear unsuitable for the South and Southeast Asia populations. However, continental Southeast Asia and northern Australia appear suitable for the Southern Hemisphere population.

**Figure 2.**
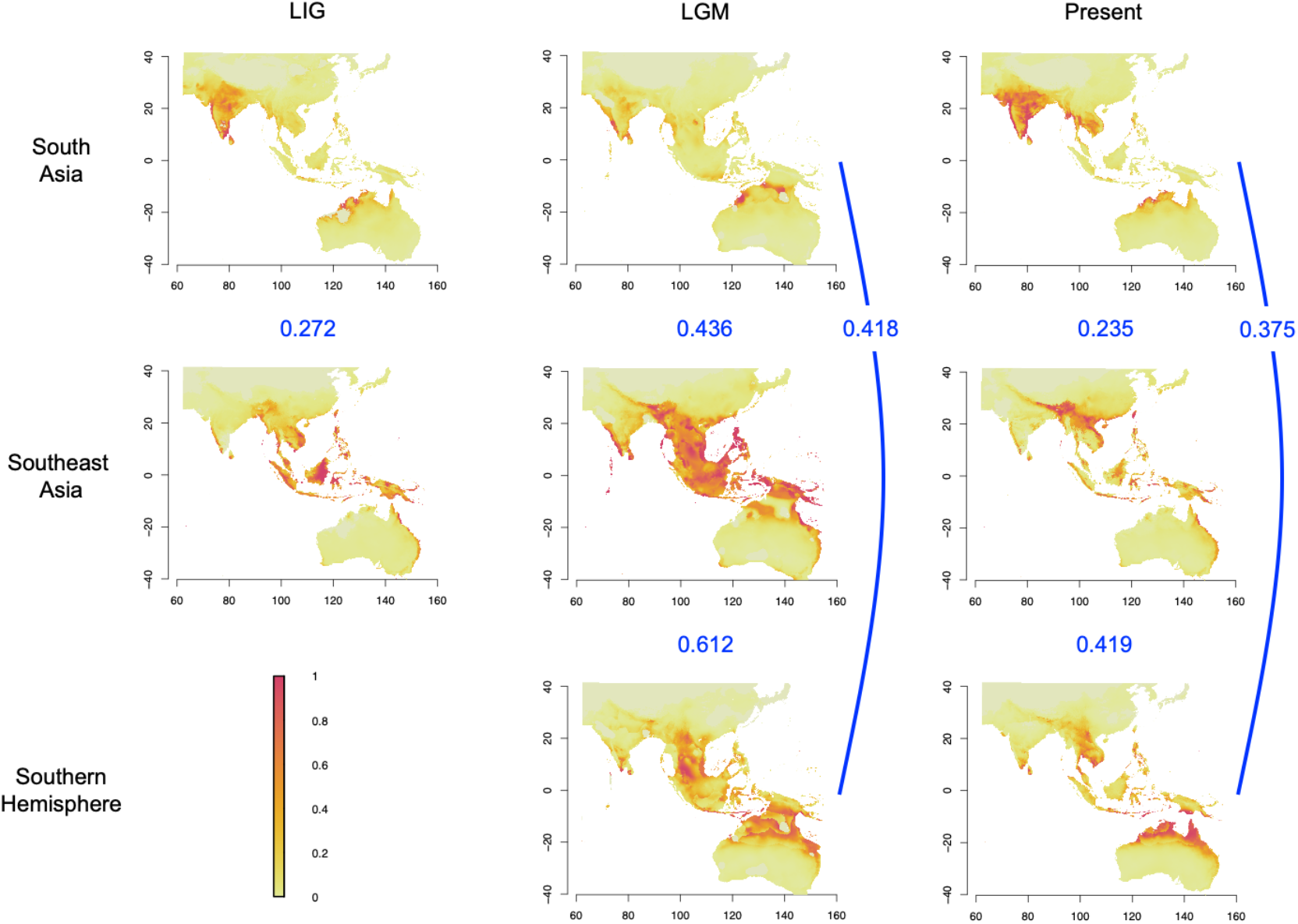
Ecological niche modeling of suitable habitats for the wild mungbean populations in South Asia, Southeast Asia, and Southern Hemisphere from the past to present. Colors represent niche suitability. The three rows (South Asia, Southeast Asia, and Southern Hemisphere) represent niche models constructed from the presence data of wild mungbean from these regions and projected to the whole geographical extent. Data of Last Interglacial period (120-140 kya, LIG) and the Last Glacial Maximum (22 kya, LGM) were based on the circulation model of MIROC-ESM. The numbers in blue between pairs of modeled distribution indicate the Schoner’D values, where higher values represent higher niche similarity.

### Selective sweeps in mungbean

We used three approaches to investigate selection footprints across the genome. Potentially selective signals across the genomes of wild and domesticated mungbeans were detected by identifying the regions with the top 1% value in the composite likelihood ratio (CLR) analysis (Figure 3A-B). There were 429 genes detected across the wild genome, and the gene ontology (GO) analysis showed that these genes were significantly enriched in the function of the protein binding and metabolic process (Supplementary Fig. S7A). Comparing our selection scan results to previous QTL mapping explicitly focusing on the divergence between wild and cultivar accessions (Isemura, et al. 2012), the wild accessions’ alleles of these QTL generally have more pods, a higher proportion of shattering pods, and smaller seed size.

**Figure 3.**
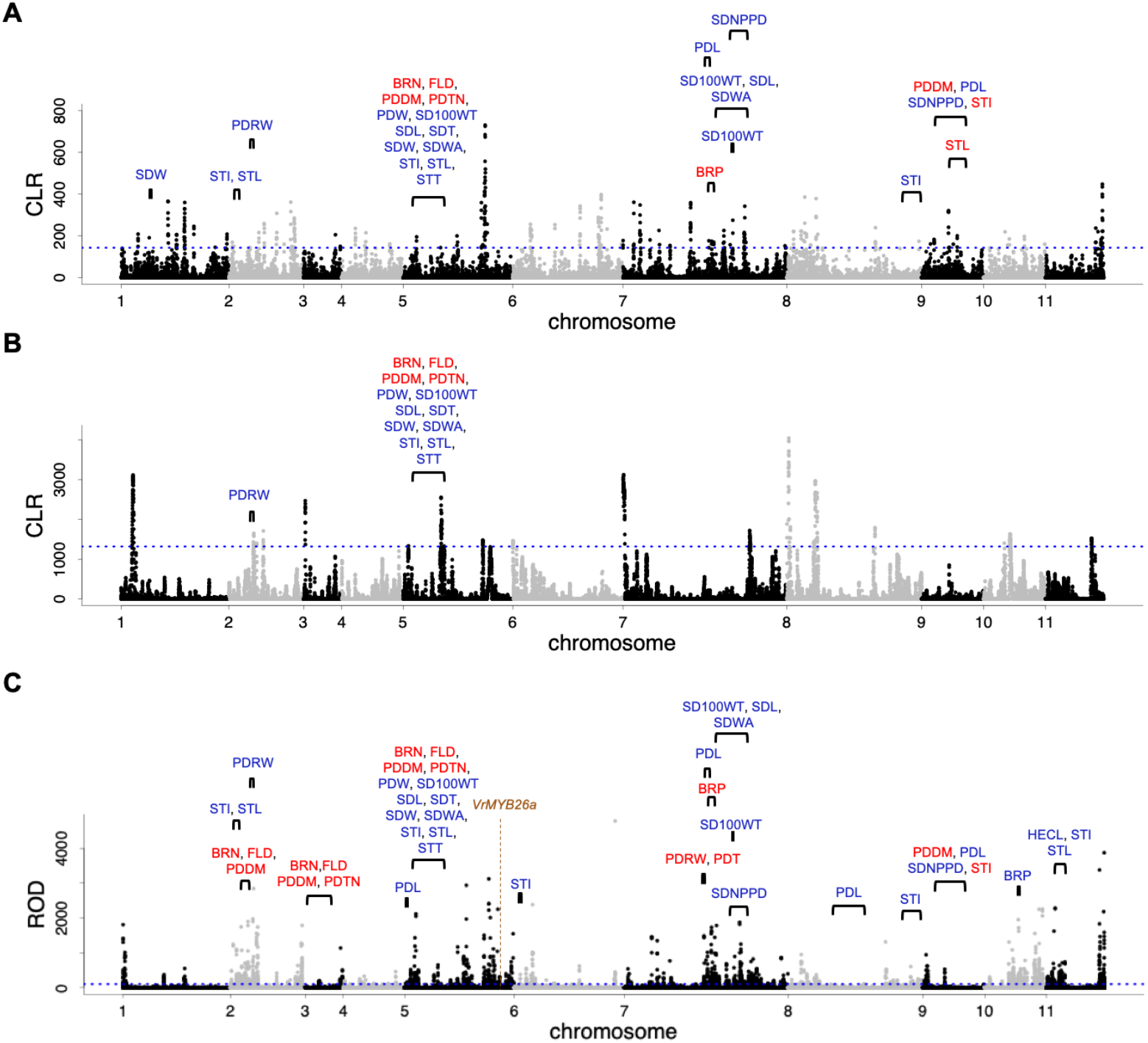
Signatures of selection across the mungbean genome. Composite likelihood ratio (CLR) values across the genomes of the (A) wild and (B) domesticated mungbean. (C) Selective sweeps during domestication inferred from reduction of diversity (ROD) values. To improve the accuracy, only the QTL with the interval < 15 Mb and overlapped with selection signals were labeled. The red and blue texts indicate wild or cultivar alleles have higher trait values in the QTL, respectively. The blue horizontal dashed lines indicate the cut-off (top 1%) of candidate selective signals. One of the candidate domestication genes, *VrMYB26a*, was labeled with brown color. (Abbreviation of traits-BRN: branch number, BRP: position of the first branch, FLD: days to first flower, HECL: hypocotyl plus epicotyl length, PDDM: days to maturity of the first pod, PDL: pod length, PDRW: rate of shattered pods, PDT: number of twist of the shattered pod, PDTN: total pod number, PDW: pod width, SD100WT: 100 seed-weight, SDL: seed length, SDNPPD: number of seeds per pod, SDT: seed thickness, SDW: seed width, SDWA: seed water absorption, STI: stem internode length, STL: stem length, STT: stem thickness).

For cultivated mungbean, the selected regions across the cultivar genome colocalized with the QTL where the cultivar alleles conferred lower pod dehiscence and fewer pods. There were 420 candidate genes in selected regions, and the GO terms were enriched in carbohydrate binding, response to biotic stimulus, and regulation of nitrogen compound metabolic process (Supplementary Fig. S7B). Notably, legumes are able to form a symbiotic relationship with rhizobia, a unique characteristic involving in nitrogen fixation process(Janczarek, et al. 2015). Thus, enrichment results suggested the important role of mungbean cultivars for soil fertility. This is supported by the first written record of mungbean in an ancient Chinese agricultural literature (齊民要術, Qimin Yaoshu, 544 AD), which emphasized its value as green manure. Other ancient Chinese sources also repeatedly mentioned using mungbean to restore soil fertility in the rotation system (Chen 1980).

A total of 1,232 genes have potential signatures of selective sweeps from the reduction of diversity (ROD) method comparing cultivars relative to wild accessions (Figure 3C). These candidate genes were enriched in GO terms such as catalytic activity, DNA binding, developmental growth, and response to stimulus and chemicals (Supplementary Fig. S7C). Among them, *INVA* is one of the candidate genes responsible for seed size in durum wheat (Mangini, et al. 2021). *ARG1* is the ortholog of the omega-3 fatty acid desaturase gene (*FAD3*), which influences the oil composition in soybean seeds (Bilyeu, et al. 2005; Pham, et al. 2012). *BG7S-2* is likely responsible for basic 7S globulin, one of the storage proteins in mungbean seeds (Mendoza, et al. 2001; Hirano 2021). The high-ROD regions colocalized with the QTL regulating phenology, pod and seed enlargement, pod twisting and dehiscence, and yield. Overall, these results suggested that the wild mungbean was likely under natural selection for dispersal: the wild alleles of these QTL generally have more pods, a higher proportion of shattering pods, and smaller seed size. On the other hand, the cultivars were selected for pod non-shattering as well as pod and seed gigantism, with fewer pods as a potential trade-off.

### *VrMYB26a* associated with the domestication trait pod non-shattering

In legumes, loss of pod twisting is one of the essential traits during domestication (Fuller and Allaby 2018). Among all the putative domestication genes, *MYB26*, which is involved in pod shattering in *Phaseolus vulgaris*, *Vigna angularis*, and *V. unguiculata* (Takahashi, et al. 2020; Di Vittori, et al. 2021), was detected to be under positive selection in the cultivars (Figure 3C). The thickness of the lignified sclerenchyma layer on the pod walls strongly correlates with the coiling of pod walls (Takahashi, et al. 2020; Parker, et al. 2021). Like other legume species, mungbean cultivars have a thinner lignified layer than wild accessions (Figure 4A). Mungbean has two *MYB26* copies, one on chromosome 5 (*VrMYB26a*) and the other on chromosome 9 (*VrMYB26b*). According to the phylogenetic relationship with the other legume homologs, *VrMYB26a* was clustered with the *MYB26* orthologs associated with pod shattering in *P. vulgaris*, *V. angularis*, and *V. unguiculata* (Clade A), and the other clade contained *VrMYB26b* (Clade B) (Supplementary Fig. S8A). The genes in Clade A showed lower *d_N_*/*d_S_* than those in Clade B (Supplementary Fig. S8B), indicating a stronger selection constraint in the former clade. Despite larger difference of allele frequency of three non-synonymous SNPs between wild and cultivars in *VrMYB26a* coding region, the function of its encoding protein is probably not changed due to the similar property of amino acids. (Supplementary Fig. S9). We inferred whether the differential expression, instead of amino acid changes, results in non-twisting pods in cultivars.

**Figure 4.**
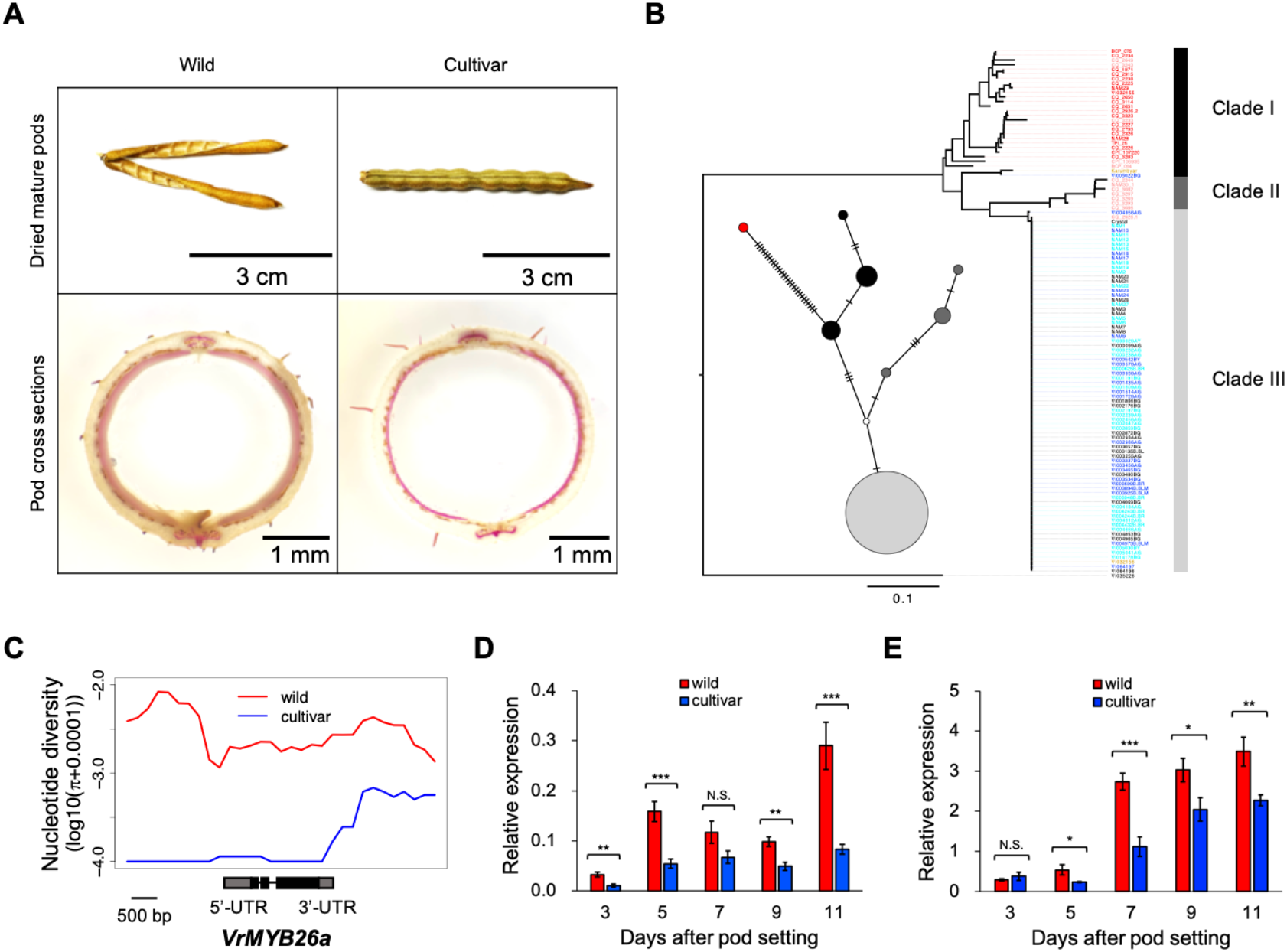
*VrMYB26a* associated with pod twisting in mungbean. (A) Mature pods and pod cross-sections from the wild (NAM30) and cultivated (VI004853) mungbean accessions. The cross-section’s lignin was stained in red. (B) Neighbor-joining tree of *VrMYB26a* promoter region (upstream 2kb) of 114 mungbean accessions and one *V. mungo* as outgroup. Accession names are labeled with colors based on population structure (K = 5). Also shown is the haplotype network based on 61 SNPs in the same region, and the outgroup was labeled with a red dot. Haplotypes were colored based on the clades identified from the tree. The sizes of circles correspond to the number of individuals in this haplotype. (C) Patterns of nucleotide diversity in sliding windows (window size, 1 kb; step size, 0.2 kb) of the wild and cultivar groups from the 2-kb upstream of *VrMYB26a* to its 2-kb downstream regions. (D) Expression levels of *VrMYB26a* during pod development. (E) Expression levels of *VrCAD4* during pod development. In (D) and (E), three accessions of wild and cultivar groups were used, respectively. The data are presented as mean ± standard error (N.S. indicates non-significant; *, **, and *** indicate *p* value < 0.05, 0.01 and 0.001, respectively).

The neighbor-joining tree and haplotype network of the *VrMYB26a* promoter region (upstream 2kb) supported the near-fixation of the derived allele in cultivars (Figure 4B). Together with the apparent reduction of nucleotide diversity at the upstream of *VrMYB26a* in cultivars (Figure 4C), these lines of evidence suggested that the expression pattern of *VrMYB26a* was the likely target of selection for non-twisting pods. We then evaluated the gene expression of *VrMYB26a* during pod development. The wild accessions showed significantly higher expression of *VrMYB26a* than cultivars on average, except on the 7th day after pod setting (DAP) (Figure 4D). Similarly, the expression of *VrCAD4*, a critical downstream gene involved in lignin biosynthesis (Xie, et al. 2018), was significantly higher in wild accessions than cultivars, except on the 3th DAP (Figure 4E). This suggested the lower expression of *VrMYB26a* reduced expression of downstream lignin biosynthesis gene, which contributed to non-twisting pods with thinner sclerenchyma layer. The gene responsible for pod non-shattering, an important trait in mungbean domestication, likely experienced the classic pattern of a hard selective sweep, where most cultivars share the same derived allele with low variation.

### *VrDet1* associated with the improvement trait stem determinacy

Plant architecture, associated with seed yield and environmental adaptation, is often the target of artificial selection during the improvement phase (Huyghe 1998). Like other legume species, wild mungbeans have indeterminate growth, while many cultivars have determinate growth (Nguyen, et al. 2012; Chauhan and Williams 2018), an improved phenotype advantageous for harvesting due to the shorter reproductive periods (Nair, et al. 2020). A previous study has shown that *VrDet1* modulated determinacy (Li, et al. 2018). The phenotypic shift from indeterminate to determinate growth was caused by two single nucleotide substitutions at the -1058 and -14 sites upstream of *VrDet1*, reducing its expression. While the previous study was based on only a few wild and cultivar accessions, our comprehensive sampling identified several cultivars with the indeterminate phenotype (Figure 5A). Thus, we further investigate patterns of polymorphism in this essential gene.

**Figure 5.**
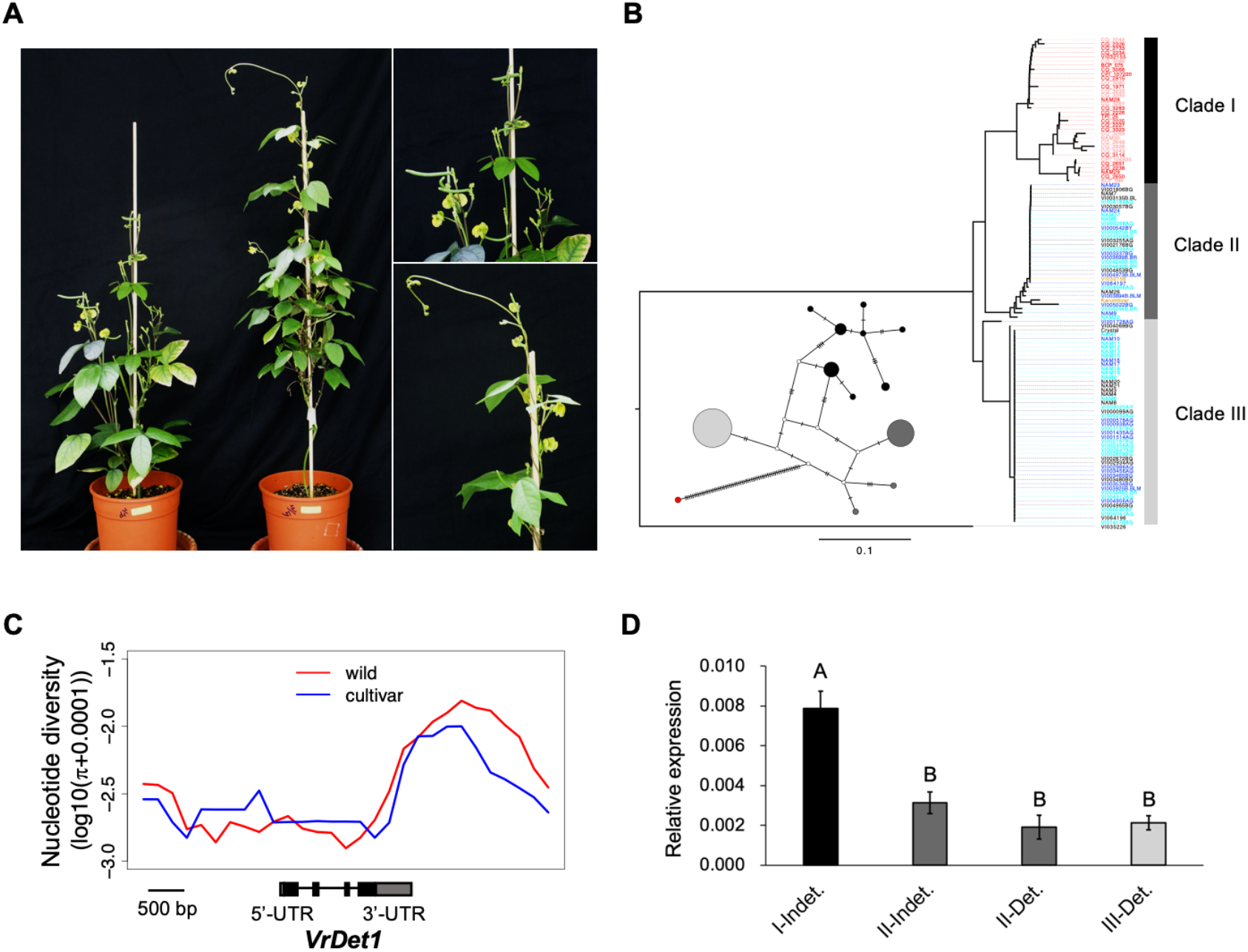
Stem determinacy and its relationship with *VrDet1*. (A) Left: Photographs of indeterminate and determinate mungbean accessions at the reproductive stage. Upper right: the enlargement of terminal flowers and pods at the main apical stem in a determinate accession. Lower right: the enlargement vegetative growth at the main apical stem in an indeterminate accession. (B) Neighbor-joining tree of the *VrDet1* promoter region (upstream 2kb) of 114 mungbean accessions and one *V. mungo* as an outgroup. Accession names are labeled with colors based on population structure (K = 5). Also shown is the haplotype network based on 85 SNPs in the same region, and the outgroup was labeled with a red dot. Haplotypes were colored based on the clades identified from the tree. The sizes of circles correspond to the number of individuals in this haplotype. (C) Patterns of nucleotide diversity in sliding windows (window size, 1 kb; step size, 0.2 kb) of the wild and cultivar groups from the 2-kb upstream of *VrDet1* to its 2-kb downstream regions. (D) Expression levels of *VrDet1* in the four groups. Colors represent the clades in (B). The data are presented as mean ± standard error (*p* < 0.001. Different letters indicate significant differences from Tukey’s HSD-test).

Non-synonymous SNPs were not found in the *VrDet1* coding region among cultivars, suggesting that the polymorphism of determinacy in cultivars was not associated with amino acid changes in this gene (Supplementary Fig. S10). The only two non-synonymous SNPs across all of our samples also have low allele frequency differences between cultivar and wild accessions (Supplementary Fig. S10). The neighbor-joining tree of the putative promoter region of *VrDet1* (upstream 2 kb) showed three distinct clades. All Australian wild (SubAU) accessions were clustered in Clade I, two Asian wild (SubAS) accessions clustered in Clade II, and the cultivars clustered in Clade II and III (Figure 5B). The haplotype of the two causal SNPs in Clade I was T^-14^/T^-1058^, and that in Clade III was A^-14^/C^-1058^, respectively corresponding to the typical wild and cultivar forms observed in the previous study (Li, et al. 2018). Interestingly, Clade II represents a novel haplotype, T^-14^/C^-1058^, with the first site corresponding to the wild but the second corresponding to the cultivar alleles. Among accessions with the Clade II haplotype, the two SubAS accessions and eight cultivars were indeterminate, and 22 out of 30 cultivars showed determinate growth. Almost all cultivars in Clade III showed determinate growth.

To test whether Clade II (T^-14^/C^-1058^) originated from a recent recombination event between the wild Clade I (T^-14^/T^-1058^) and the “typical cultivar” Clade III (A^-14^/C^-1058^) haplotypes, in this upstream 2 kb region we identified nine diagnostic SNPs with high allele frequency differences between Clade I and III. We used these SNP sites to polarize the allele frequencies in the newly identified Clade II. Instead of the typical patterns of recent recombinant haplotype, where one side of the Clade-II haplotype would conform to Clade-I and the other correspond to Clade-III major alleles, the allele frequencies of Clade II zigzagged among these SNPs (Supplementary Fig. S11). This suggests that Clade II (T^-14^/C^-1058^) is an ancient haplotype, which is also supported by the haplotype network where haplotypes in both Clade II and Clade III are closer to the root (Figure 5B). Since the cultivars contained two haplogroups of intermediate frequencies, no obvious lack of variation was observed in cultivars compared to wild accessions (Figure 5C). On the other hand, many accessions within Clade II or Clade III possess identical sequences, suggesting their independent rapid frequency increase (Figure 5B).

The gene expression level of *VrDet1* in Clade II and III was significantly lower than that in Clade I (Figure 5D), consistent with their determinate phenotype. Notably, the indeterminate accessions in Clade II did not show significantly higher expression than the determinate accessions in Clade II or Clade III, suggesting *VrDet1* expression might not be the only factor affecting stem determinacy. The results suggested that Clade II (T^-14^/C^-1058^) may have weaker effects than Clade III (A^-14^/C^-1058^) and have incomplete penetrance, sometimes allowing other genes to cause the ancestral indeterminate phenotype. This is consistent with Clade II containing one wild (T^-14^) and one cultivar (C^-1058^) allele in the two SNPs previously shown to affect gene expression (Li, et al. 2018).

Taken together, our results suggested that the evolution of the improvement trait, stem determinacy, appears to be more complex than the previously suggested simple model of hard selective sweep from the "typical cultivar" Clade III haplotype (Li, et al. 2018). Instead, the evolution might result from soft sweeps of two ancient haplotypes (Clade II T^-14^/C^-1058^ and Clade III A^-14^/C^-1058^) reducing the expression of *VrDet1*, with Clade II showing incomplete penetrance allowing the indeterminate phenotype under certain environmental or genomic contexts.

## Discussion

In this study, we used whole-genome sequencing to investigate the history of *Vigna radiata* before, during, and after domestication. Our results suggest *Vigna radiata* originated in Asia, and the wild population of Southeast Asia likely migrated to Australia during the last ice age. This is supported by the geographical proximity and niche similarity between Southeast Asia and Australia. Domestication probably happened at 9 kya in Asia, after which the cultivars had high selection pressure during domestication for loss of pod shattering to prevent yield loss. Subsequently, some cultivars with determinate shoot growth were selected during improvement. Given the different nature of selection during the domestication and improvement phases, we showed that the candidate genes for these two traits exhibit distinct signatures of selection.

### The evolution of *V. radiata*

We proposed a hypothesis of the evolution of *V. radiata*: wild mungbean originated in Asia and migrated into Australia at about 50 kya. This coincides with the initial occupation of Australia by humans (Tobler, et al. 2017), and interestingly, it has been recorded that the tuberous roots of wild mungbean were consumed by Aboriginal Australians (Lawn, et al. 1988). Whether the immigrations of mungbean and human into Australia are associated remains to be further investigated. Compared to their Asian counterparts, wild Australian mungbean evolved smaller seeds (Supplementary Table S3) (Lawn and Rebetzke 2006), likely as an adaptive strategy for dispersal. While wild populations both have twisting pods, the domestication starting at about 9kya further selected for pod non-twisting (Figure 6). Traditional mungbean cultivars were indeterminate (Nair, et al. 2020), and determinate cultivars were further selected during improvement. The phenomenon of two-step artificial selection also occurred in other crops. For example, in rice, seed dormancy and seed shattering were under artificial selection during domestication, while plant height and flowering time were modified locally during improvement (Liu, et al. 2018). The two-step selection in similar traits also occurred in wheat (Roucou, et al. 2018). In tomatoes, yield-related traits were first selected during domestication, and traits of flavor and color were selected in cultivars during improvement (Schouten, et al. 2019).

**Figure 6.**
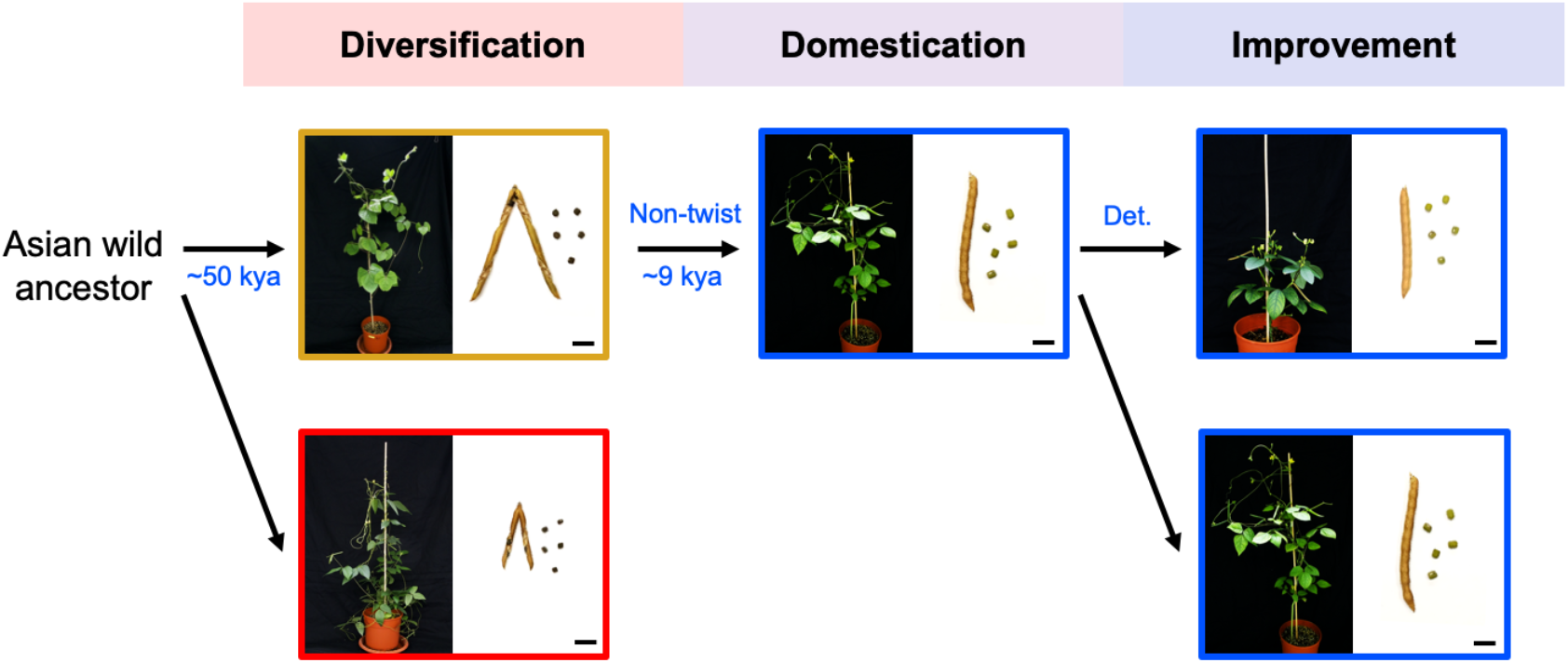
The hypothesis of mungbean evolution. A wild progenitor of *V. radiata* experienced species diversification, resulting in the divergence of *V. radiata* var. *sublobata* into SubAS (goldenrod box) and SubAU (red box) in about 50 kya. SubAS was further selected for pod non-twisting, becoming *V. radiata* var. *radiata* (blue boxes) during domestication (about 9 kya). Afterwards, some indeterminate cultivars are improved into determinate forms. Bar = 1 cm.

### Different selection forces resulting in distinct patterns of genetic variation

In legumes, mutations of genes for pod shattering reduce the torsion of pod walls or strengthen the sutures, contributing to non-shattering pods (Parker, et al. 2021). Our result showed that the lower expression of *VrMYB26a* in mungbean cultivars is associated with the thinner sclerenchyma layer in non-twisting pods. In *Vigna angularis* and *V. unguiculata,* it has been shown that mutations creating truncated protein of *MYB26* are likely responsible for non-twisting pods in cultivars (Takahashi, et al. 2020). In *Phaseolus vulgaris*, down-regulation of *PvMYB26* contributed to indehiscent pods (Di Vittori, et al. 2021). These findings reflects Vavilov’s Law of Homologous Series in Variation, which stated that parallel evolution in closely-related species often involve genetic changes in homologous genes due to their conserved functions (Vavilov 1922, 1951). In addition, although archeological evidences suggested two possible domestication origins of mungbean from northern and southern India (Fuller and Harvey 2006), only one haplotype was observed in *VrMYB26a* promoter region across all cultivars. We suggested that mungbean likely went through only one domestication event for non-twisting pods. Consistent with the general trend in domestication genes under strong directional selection, we found evidences of hard selective sweep in *VrMYB26a*.

As for stem determinacy, indeterminate and determinate growth widely exists in cultivars (Eshed and Lippman 2019). Indeterminate crops have relatively high yields because they provide extended harvest periods and unconstrained canopy development (Chauhan and Williams 2018). Determinate types have short plant height, lodging resistance, and uniform maturity properties (Lawn 1989; Kato, et al. 2019). These two growth habits have different performances under different climate regions or agricultural practices. In the northern regions of America and China, indeterminate soybean varieties are commonly used due to their greater yield under lower temperatures and limited rainfall (Specht, et al. 2014; Kato, et al. 2015; Liu, et al. 2015). In contrast, determinate varieties dominate southern regions (Kilgore-Norquest and Sneller 2000). In mungbean, subsistence farmers prefer cultivating indeterminate varieties for multiple harvesting (Hewavitharane, et al. 2010; Depenbusch, et al. 2021), whereas determinate varieties were commonly used in intensive farming (Nair, et al. 2020). These two types of mungbean cultivars have existed in China since at least the 16^th^ century, as recorded in The Compendium of Materia Medica (本草綱目 Ben Cao Gang Mu, 1578 AD) (Li 2016) and Chinese encyclopedia of technology (天工開物 Tian Gong Kai Wu, 1637 AD) (Song, et al. 1997). The previous study identified alleles (A^-14^ and C^-1058^) in two SNPs of *VrDet1* responsible for determinate growth in mungbean (Li, et al. 2018). We found that two ancient haplotypes (Clade II T^-14^/C^-1058^ and Clade III A^-14^/C^-1058^) reduced *VrDet1* expression, suggesting these two haplotypes were selected when determinate varieties were preferred during improvement and consistent with soft selective sweeps (Garud, et al. 2015; Harris, et al. 2018). In soybean, multiple haplotypes of *GmTFL1* associated with determinate growth in soybean were also observed (Liu, et al. 2015). In the cross between two accessions with Clade I and Clade III haplotypes, Li et al. showed perfect co-segregation between determinacy traits and SNP markers in the promoter region (Li, et al. 2018). This suggests the complete penetrance and strong effect of Clade III (A^-14^/C^-1058^). It’s worth noting that some cultivars with Clade II haplotype (T^-14^/C^-1058^) displayed an indeterminate growth, suggesting this haplotype may have incomplete penetrance allowing the generation of indeterminate phenotype under specific environmental or genomic contexts (Pierce 2012). Under specific environments or agricultural practices where the indeterminate phenotype was preferred, the Clade II haplotype may be favored. In brief, our results showed that multiple adaptive haplotypes of *VrDet1* were selected in mungbean cultivars, resulting in a pattern distinct from the typical signatures of hard selective sweep.

## Conclusion

In this work, we investigated the evolutionary history of *Vigna radiata* using genomic approaches. Our comparison of phylogenetic tree, haplotype network, and signature of selection at *VrMYB26a and VrDet1* loci elucidated distinct genetic architecture behind different phases of artificial selection. For a trait and the relevant gene under selection during the domestication phase, pod twisting and *VrMYB26a* in our case, a hard selective sweep was detected with the near-complete fixation of one derived haplotype in cultivars. For a trait and the relevant gene during the improvement phase, determinacy and *VrDet1* in our case, phenotypic polymorphisms and genetically diverse haplotypes at the locus were observed in cultivars. With whole-genome resequencing of diverse wild and cultivars, we clarify the demographic history of mungbean and revealed distinct patterns of genes affected by artificial selection during crop domestication and improvement phases.

## Materials and Methods

### Plant materials, library construction, and sequencing

114 *V. radiata* accessions and an outgroup, *V. mungo*, were obtained from two genebanks: the World Vegetable Center in Taiwan and the Australian Grains Genebank (AGG). The passport data from the genebanks provided the taxonomy and country of origin for each accession (Supplementary Table S2). The genomic DNA was extracted from leaves with the DNeasy Plant Mini Kit (QIAGEN). DNA libraries were constructed using NEBNext® Ultra™ II DNA Library Prep Kit for Illumina (E7645), and 150-bp paired-end sequencing was completed using Illumina Hiseq X Ten.

### Variant calling

Raw sequencing reads were trimmed with SolexaQA++ v3.1.7.1 (Cox, et al. 2010). Then adaptor sequences were removed with cutadapt v1.14 (Martin 2011). The cleaned reads were mapped to the mungbean cultivar (VC1973A) reference genome version 6 (Vradiata_ver6) (Kang, et al. 2014) with Burrows-Wheeler aligner v0.7.15 (Li and Durbin 2009). Duplicated reads were marked with Picard v2.9.0-1 (http://broadinstitute.github.io/picard/). Single-nucleotide polymorphisms (SNPs) were called with GATK v3.7 following the GATK Best Practice (McKenna, et al. 2010; Van der Auwera, et al. 2013). We used vcftools v0.1.13 (Danecek, et al. 2011) with following parameters “--min-alleles 2 --max-alleles 2 --remove-indels --max-missing 0.9 --minQ 30” to filter and retain bi-allelic SNPs. For LD decay and fixation index (*F_ST_*) analyses, SNPs with minor allele frequency (MAF) < 0.05 were further removed.

### Gene prediction and annotation

Given that the public gene annotation of *V. radiata* contains only 29,006 genes, which are much less than closely related species (Kang, et al. 2014), we re-annotated the reference genome comprehensively. The whole Vradiata_ver6 genome without masking repeats was annotated ab initio with Augustus v3.3.2 using -species Arabidopsis option (Stanke, et al. 2006). For RNAseq evidence, we downloaded RNA sequencing reads from seeds, pods, flowers, and one-week-old and four-week-old whole plants (Chen, et al. 2015; Liu, et al. 2016). These RNA reads were mapped onto the Vradiata_ver6 genome with HISAT v2.1.0 (Kim, et al. 2015) and assembled and merged by StringTie v1.3.5 (Pertea, et al. 2015). We then blasted the assembled transcripts using blastp v2.8.2 (Camacho, et al. 2009) on the UniProt database (https://www.uniprot.org/) and predicted Open Reading Frames using TransDecoder v5.5.0 (Haas, et al. 2013). For protein evidence, the *Glycine max* protein sequences (Wm82.a2.v1) (https://soybase.org/) were used as reference protein evidence and aligned to the Vradiata_ver6 genome with exonerate v2.4(Slater and Birney 2005). All these evidences were submitted to Evidencemodeler v1.1.1 (Haas, et al. 2008) to identify consensus gene models. The weight of evidence was five except for ab initio evidence, which was one. Finally, the consensus gene models were blasted in the databases of TRAPID (http://bioinformatics.psb.ugent.be/trapid_02/), eggNOG-mapper (Huerta-Cepas, et al. 2017), and plaza (Van Bel, et al. 2018) for functional annotation.

### Niche modeling

We used the occurrence data of *V. radiata* var. *sublobata* based on the Global Biodiversity Information Facility (GBIF; https://www.gbif.org/) and also the collection sites of the wild samples from the Australian Grains Genebank (AGG) and National Bureau of Plant Genetic Resources (NBPGR) to identify the niche of the wild mungbean. The species distribution ranged from 32.927146° N, 66.501760° E to 27.820616° S, 155.770869° E. To avoid the over-represented occurrence in a single geographic grid, we removed the overlapped occurrence in the combined data, resulting in 22, 26, and 39 samples in South Asia, Southeast Asia, and Southern Hemisphere, respectively (Supplementary Table S4). Nineteen bioclimatic variables, including the current data (1960-1990) and the paleoclimatic data, were downloaded in the highest resolution from WorldClim 1.4 version (https://www.worldclim.org/data/v1.4/worldclim14.html). The paleoclimatic data refers to the periods of the Last Glacial Maximum (LGM: 22,000 years ago) under the MIROC-ESM and CCSM4 models, and the Last interglacial (LIG: 120,000-140,000 years ago) under the MIROC-ESM model. After removing the highly correlated bioclimatic variables (Pearson’s correlation), we kept six bioclimatic variables in the niche modeling, including BIO1, BIO2, BIO3, BIO12, BIO15, and BIO18, after removing the highly correlated bioclimatic variables (Pearson’s correlation coefficients > 0.8). The niche modeling and projection were performed with MaxEnt v3.4.3 using default settings (Phillips, et al. 2006).

### Population genomics

We inferred population structure with ADMIXTURE v1.3 (Alexander, et al. 2009). An accession was defined as admixed if none of its single ancestral components was more than 0.7. The neighbor-joining tree was constructed with TASSEL v5.0 (Bradbury, et al. 2007) and plotted with Figtree v1.4.4 (http://tree.bio.ed.ac.uk/software/figtree/). Nucleotide diversity (π) and fixation index (*F_ST_*) were calculated using vcftools v0.1.13 (Danecek, et al. 2011). Linkage disequilibrium (LD) between SNPs was calculated with PopLDdecay v3.41 (Zhang, et al. 2019). To examine the relationship among populations, we estimated the genetic distance of 115 accessions with a bi-allelic SNP dataset using Plink v1.90b4.5 (--distance) (Purcell, et al. 2007) and then constructed a phylogenetic network with the neighbor-net algorithm in SplitsTree v5.3 (Huson and Bryant 2006).

### Demographic history

We reconstructed the demographic history of *V. radiata* with SMC++ v1.15.2 (Terhorst, et al. 2017) since this algorithm accepts many samples for a population. Since such an algorithm treats two gene copies within a diploid individual as two randomly sampled haplotypes from the populations and *V. radiata* is a self-fertilizing species, we followed common practices for species like *Arabidopsis thaliana* and soybean (Alonso-Blanco, et al. 2016; Kim, et al. 2021). We generated artificial diploids by assembling haplotypes from two random accessions, except for the admixed accessions, including two wild accessions (NAM30, CPI_106935) and one cultivar (VI005022BG). In addition, the heterochromatic regions, which were defined by regions enriched in repetitive sequences in the Vradiata_ver6 genome, were excluded. We estimated the divergence time among cultivated *V. radiata* var. *radiata*, the wild group from Australia (SubAU), and the wild group from Asia (SubAS), assuming one generation per year with a mutation rate at 1 × 10^-8^ (von Wettberg, et al. 2018). We annotated potential effects of SNPs with SnpEff v4.3t (Cingolani, et al. 2012), and SNPs annotated as high or low impacts were used to generate unfolded 2D Site Frequency Spectrum (2D SFS) using *V. mungo* as the outgroup.

### Detection of selective sweeps

Admixed accessions between wild and cultivar groups were removed from the analyses of selection signals during domestication. Reduction of diversity (ROD; π_wild_/π_cultivar_) was calculated in 10-kb windows with a 1-kb step size. A composite likelihood ratio (CLR) test was performed with SweeD v4.0.0 (Pavlidis, et al. 2013) in non-overlapping 10-kb sliding windows across the genome. Quantitative trait loci (QTL) of the domestication traits were detected based on previous research (Isemura, et al. 2012) by anchoring their linked markers to the reference genome. Gene ontology (GO) enrichment of the genes under selection was estimated using TBtools v1.098 (Chen, et al. 2020).

### Phenotyping

The mungbean plants were grown in a growth chamber under 12 hours (30°C) light/ 12 hours (25°C) dark at 700 µmol m^−2^ s^−1^ of light. For stem determinacy, accession was defined as determinate when apical meristems stopped growing soon after flowering. For pod twisting, pods from wild and cultivar accessions were dried in an oven at 37°C for four days before phenotyping. The weight of 100 seeds was the average of three plants in each accession. The number of seeds per pod and pod length was the average of 30 pods (10 pods per plant, a total of 3 plants) of each accession.

The sclerenchyma tissues, which contain the lignified cell walls, were observed with phloroglucinol–HCl solution (Pomar, et al. 2002). We collected fresh mature pods for histological staining. The phloroglucinol–HCl solution was prepared by dissolving 0.2 g phloroglucinol in 20 ml ethanol and additional 20 ml 37% hydrochloric acid. The free-hand cross-sections of pods were obtained with a razor blade (Corrux) and incubated in phloroglucinol–HCl solution for one minute. We observed the stained slides under an optical dissecting microscope (SZ61; Olympus).

### RNA isolation and real-time qPCR for *MYB26* and *VrDet1*

To detect *MYB26* gene expression during pod development, we collected pods 3, 5, 7, 9, and 11 days after pod setting (DAP). Seeds were manually removed except pods at 3 and 5 DAP (seeds were too small). A total of 50 mg pod tissues per sample were ground with liquid nitrogen in a 1.5-ml microcentrifuge tube and treated with 65 °C 1.2 ml CTAB buffer (2% CTAB, 1% polyvinylpyrrolidone 40, 2 M sodium chloride, 100 mM Tris-HCl, 20 mM ethylenediaminetetraacetic acid, 2% beta-mercaptoethanol in nuclease-free water) (Wang and Stegemann 2010). The samples were then incubated in a water bath at 65 °C for 30 minutes and centrifuged at 1,6000 *g* for 15 minutes. The supernatant was transferred to two new microcentrifuge tubes and added equal volumes of UltraPure™ phenol:chloroform:isoamyl alcohol (25:24:1, Invitrogen). Afterward, the evenly-mixed samples were centrifuged at 1,6000 g for 15 minutes. The supernatant was transferred to a new microcentrifuge tube and mixed with equal volumes of 8M LiCl. Subsequently, the samples were kept in a -20 °C refrigerator for at least 2 hours and then centrifuged at 1,6000 *g* for 30 minutes to remove the supernatant. The pellet was washed with 1 ml 80% ethanol and centrifuged at 1,6000 *g* for 10 minutes. Finally, the pellet was dissolved in 50 µl nuclease-free water.

As for the *VrDet1*, we evaluated the expression using apical meristem at the V0 stage (unifoliate leaves are fully expanded, and the first trifoliate leaves just emerged) following the previous study (Li, et al. 2018). RNA was extracted using the total RNA isolation kit (Novelgene; Cat. No. RT0300). The extracted RNA solution was treated with DNAse I (NEB) and then cleaned up with the LiCl method mentioned above.

cDNA was synthesized from 1 µg of total RNA for each sample using SuperScript IV Reverse Transcriptase (Invitrogen) and then diluted at 1:20. The iQ SYBR Green Supermix (Bio-Rad) was used to perform real-time qPCR. mRNA’s relative expression was calculated using the 2^−ΔΔCT^ method (Livak and Schmittgen 2001) using the housekeeping gene *VrCYP20* (Li, et al. 2015). The intron-spanning primers were designed for the qPCR experiment (Supplementary Table S5).

### Evolutionary and association analyses

We used the coding sequence of *VrMYB26* (LOC106761638) to retrieve its homologous genes in azuki bean (*V. angularis*), cowpea (*V. unguiculata*), common bean (*Phaseolus vulgaris*), and *Arabidopsis thaliana* from public databases, including VigGS (Sakai, et al. 2016), Phytozome (Goodstein, et al. 2012), and TAIR (Huala, et al. 2001). Sequences were aligned, and the maximum-likelihood (ML) tree was constructed with 1,000 bootstrap replications using MEGA v11(Tamura, et al. 2021). In addition, we estimated pairwise *d_N_*/*d_S_* ratios in each clade according to the Yang & Nielsen (2000) method (Yang and Nielsen 2000) with PAML v4.9 (Yang 2007). We constructed neighbor-joining tree of promoter region of targeted genes with ape package in R v4.1.2 (Paradis and Schliep 2018). The haplotype network was constructed using PopART v1.7 with the TCS algorithm (Leigh and Bryant 2015). We used the SNPs without heterozygotes and missing values in the *VrDet1* and *VrMYB26a* promoter regions for the haplotype network analysis and used *V. mungo* as the outgroup.

## Acknowledgments

We thank Chia-Yu Chen, Pei-Wen Ong, and Jo-Wei Hsieh for the assistance in sample preparation. We are grateful to the support from National Taiwan University’s Computer and Information Networking Center for high-performance computing facilities as well as Technology Commons of College of Life Science for molecular biology assistance. C.-R.L. was funded by 107-2923-B-002-004-MY3 and 110-2628-B-002-027 from the Ministry of Science and Technology, Taiwan. C.-T.T. was funded by 107-2923-B-002-004-MY3 from the Ministry of Science and Technology, Taiwan. Y.-P.L. was supported by 110-2313-B-125-001-MY3 from the Ministry of Science and Technology, Taiwan. R.N. and R.S. were funded by the Australian Center for International Agricultural Research (ACIAR) through the projects on International Mungbean Improvement Network (CIM-2014-079 and CROP-2019-144) and by the strategic long-term donors to the World Vegetable Center: Republic of China (Taiwan), UK aid from the UK government, United States Agency for International Development (USAID), Germany, Thailand, Philippines, Korea, and Japan. E.B.-v.-W. was supported by USDA Multistate Hatch NE1710. M.S. was supported by the Ministry of Science and Higher Education of the Russian Federation under the strategic academic leadership program "Priority 2030" (Agreement 075-15-2021-1333 dated 30.09.2021). S.N. was supported by the Zumberge foundation.

## Author Contributions

C.-R.L. designed and supervised the study. Y.-P.L., H.-W.C., and C.-R.L. performed data analyses and wrote the manuscript with help from other authors. H.-W.C. and P.-M.Y. collected phenotypic data and conducted qPCR experiment. All authors read and approved the manuscript.

## Data Availability

The raw sequencing data for each of 114 *Vigna radiata* accessions and 1 *V. mungo* generated in this study have been submitted to the NCBI BioProject database (https://www.ncbi.nlm.nih.gov/bioproject/) under accession number PRJNA838242.

---

**Supplementary Fig. S1.**
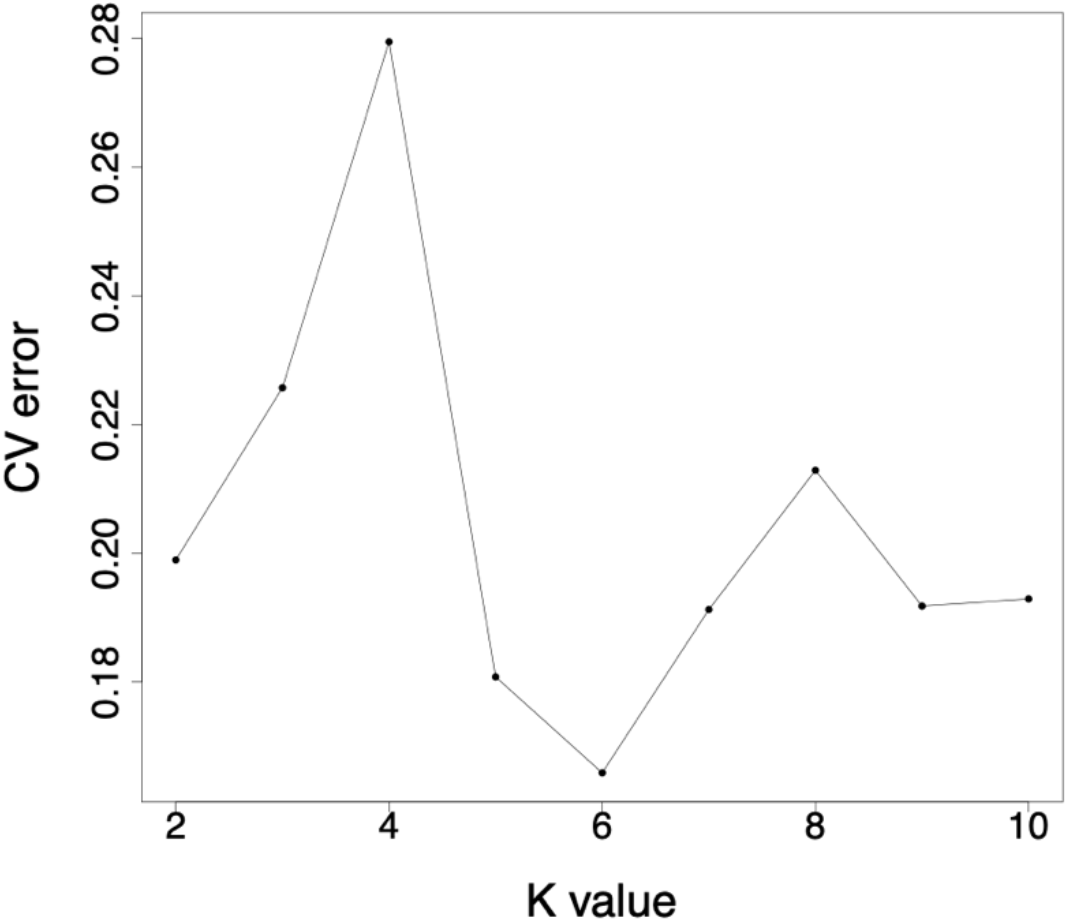
Cross-validation error of each K value.

**Supplementary Fig. S2.**
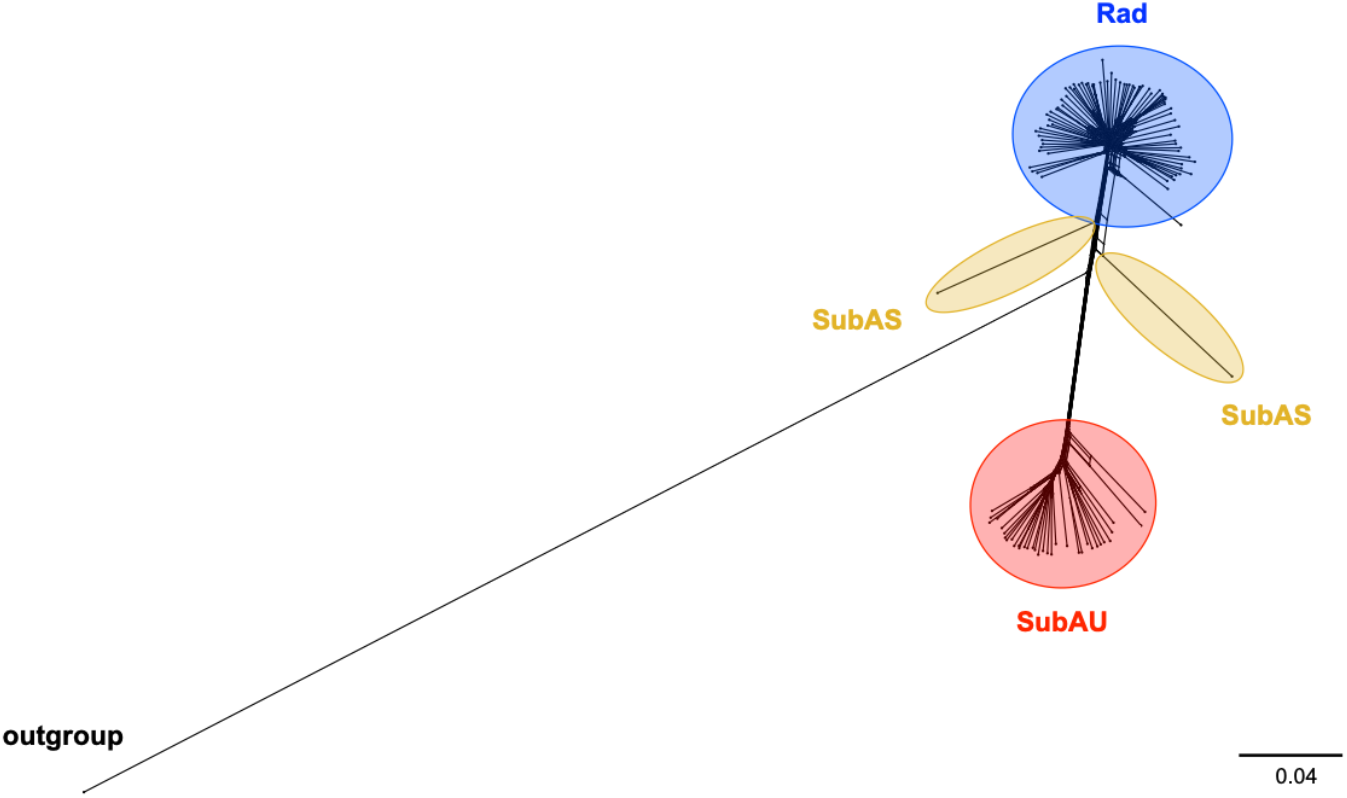
Phylogenetic network of the 114 mungbean accessions and one outgroup.

**Supplementary Fig. S3.**
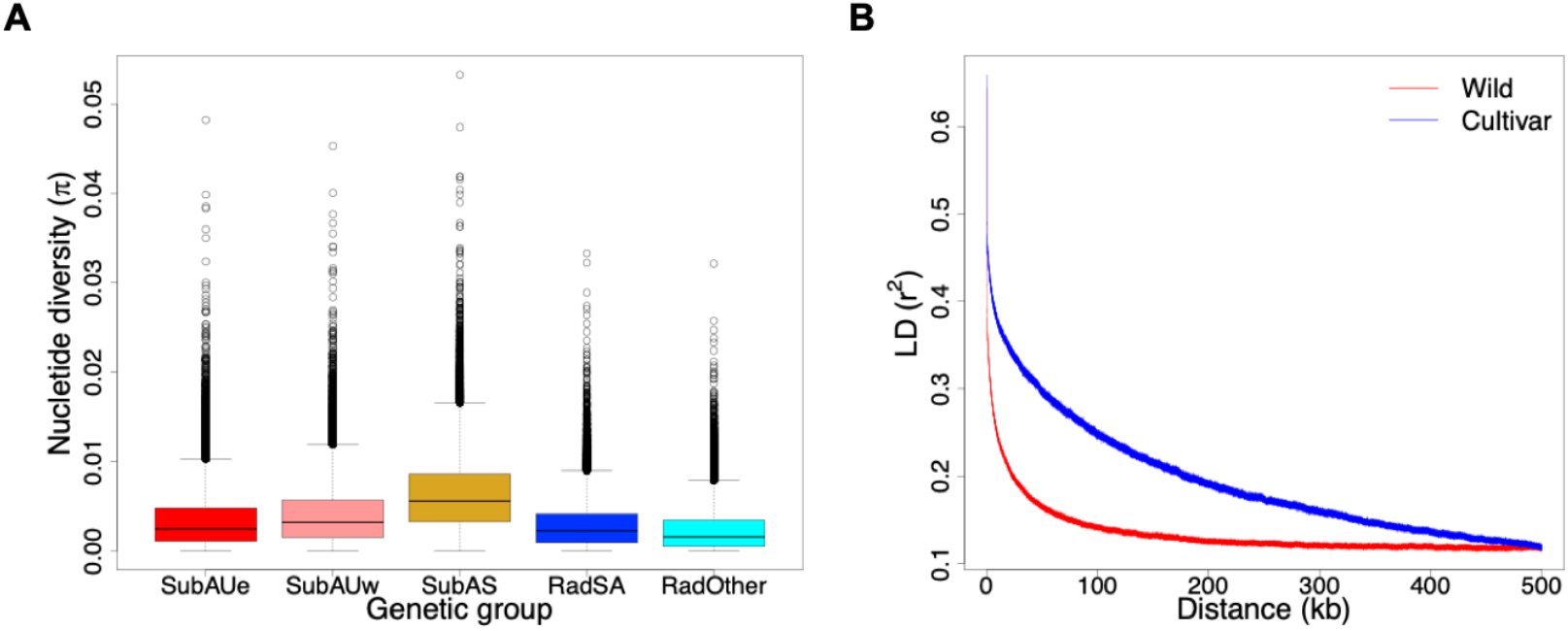
Genetic diversity of mungbean. (A) Nucleotide diversity (π) in the 10kb window of the three wild genetic groups (SubAUe, SubAUw, and SubAS) and the two cultivated genetic groups (RadSA and RadOther). The boxes indicate medians and interquartile ranges, whiskers indicate 95% values, and additional points in each boxplot represent outliers. (B) Decay of linkage disequilibrium (LD) of wild and cultivated populations.

**Supplementary Fig. S4.**
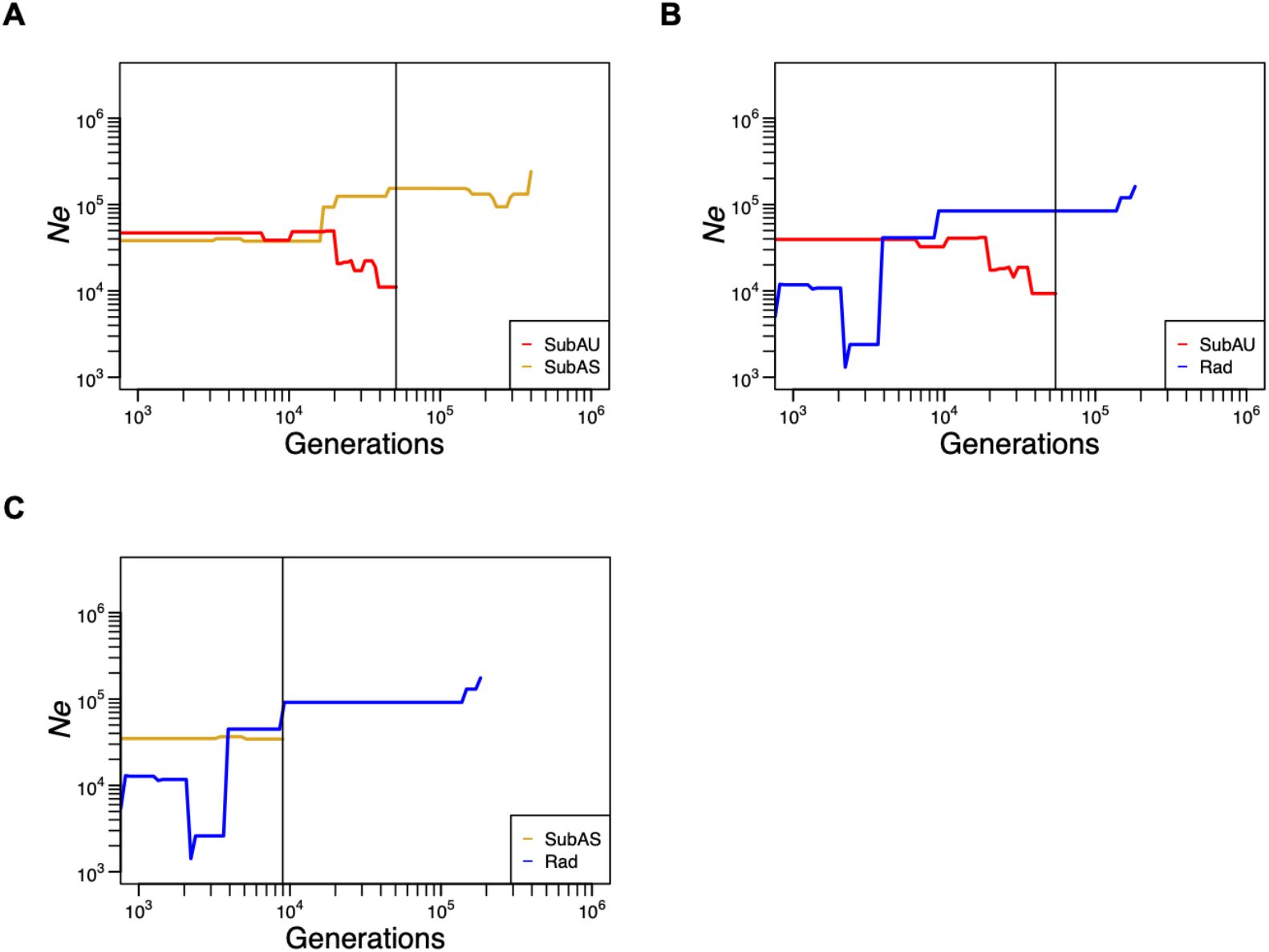
Pairwise population divergence time. (A) SubAU and SubAS (B) SubAU and Rad (C) SubAS and Rad. The vertical black lines denote the estimated divergence time.

**Supplementary Fig. S5.**
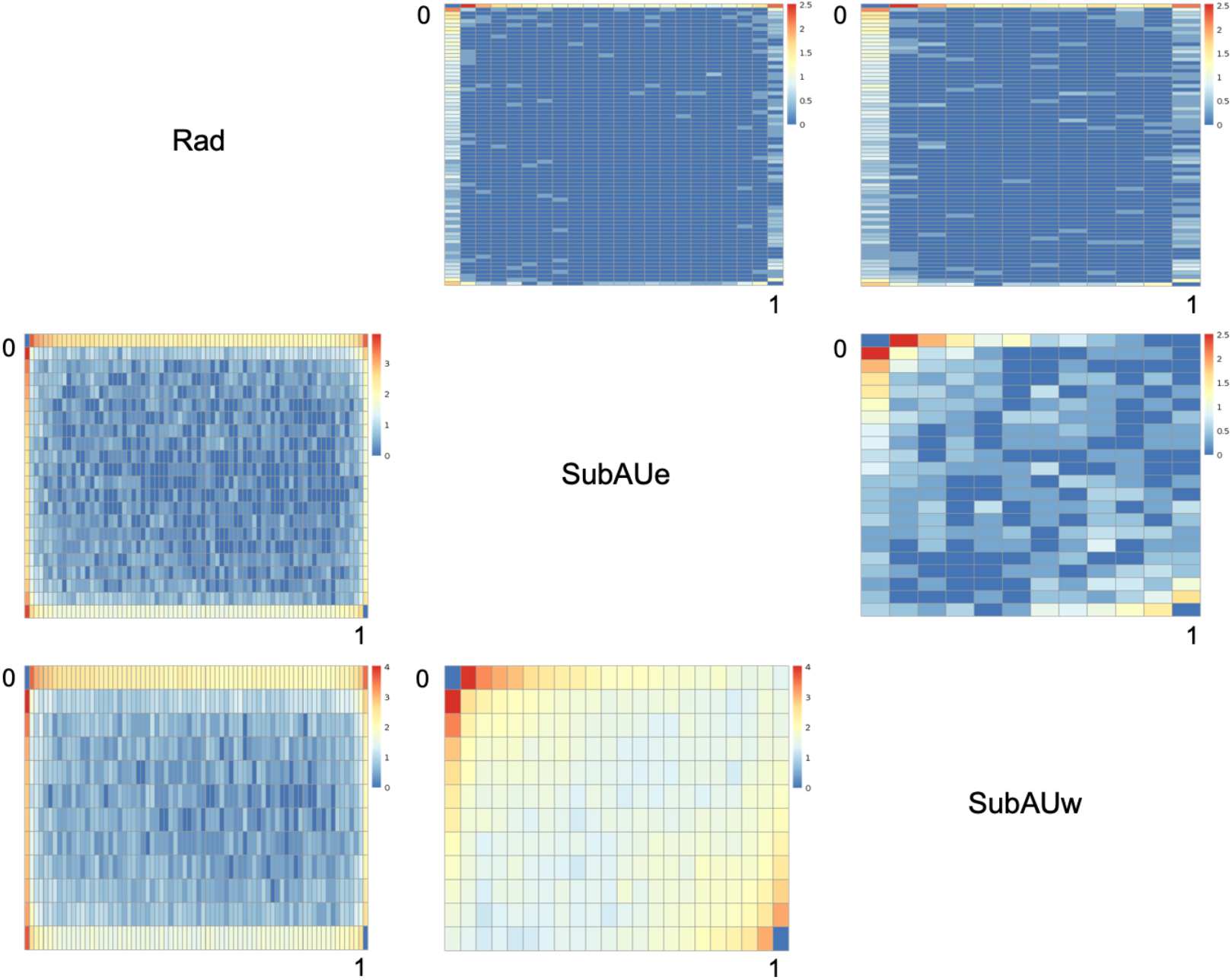
Two-dimensional site frequency spectrum (2D SFS) between Rad, SubAUe, and SubAUw groups. The upper triangle plots are 2D SFS of the SNPs with high impact estimated by SnpEff; the lower triangle plots are the 2D SFS of the 4-fold degenerate SNPs. Colors reflect the log10 number of SNPs.

**Supplementary Fig. S6.**
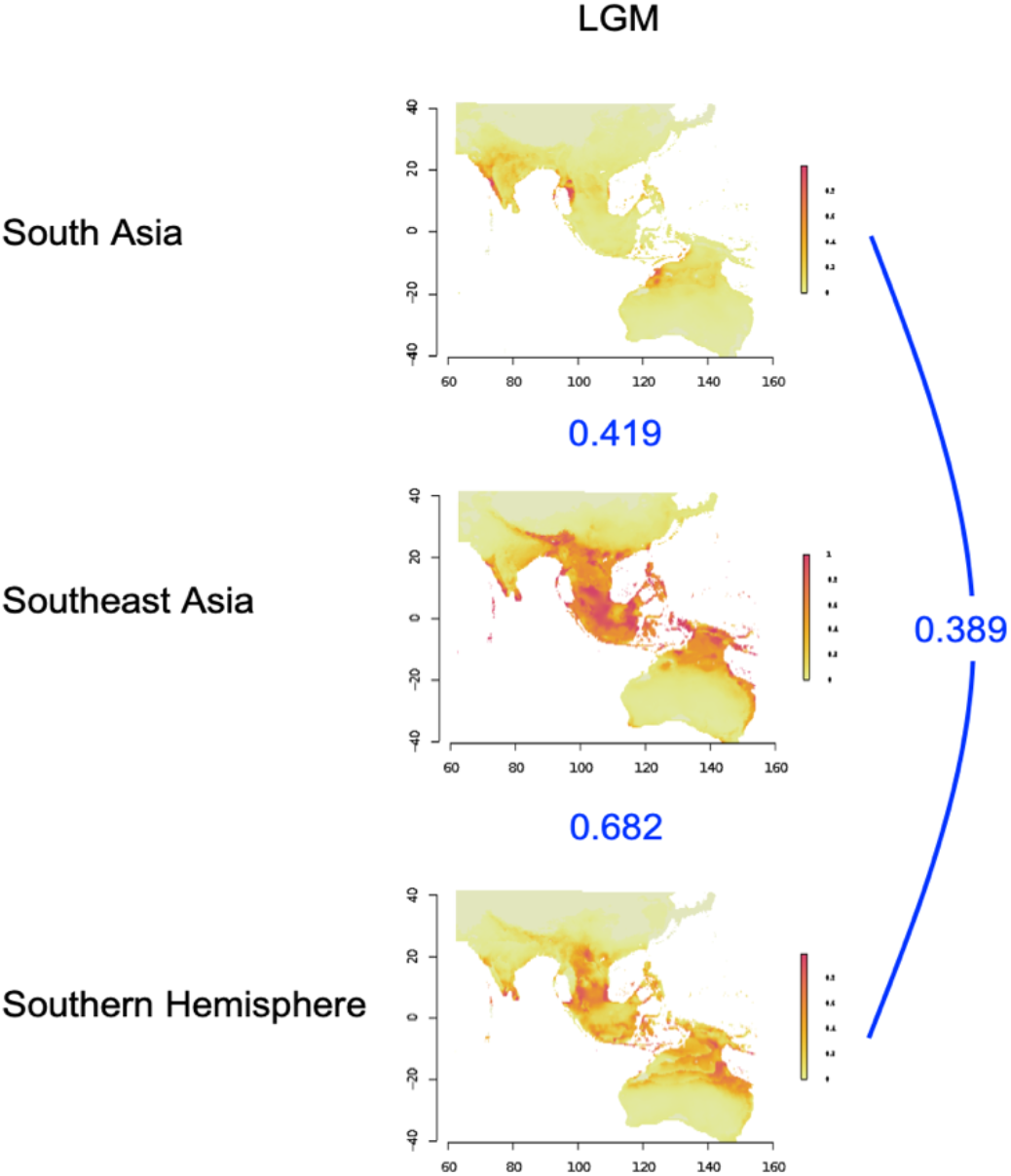
Ecological niche modeling of suitable habitats for the wild mungbean populations in South Asia, Southeast Asia, and South Hemisphere in Last Glacial Maximum (22 kya). Colors represent niche suitability. The three graphs (South Asia, Southeast Asia, and Southern Hemisphere) represent niche models constructed from the presence data of wild mungbean from these regions and projected to the whole geographical extent. Data of the Last Glacial Maximum (22 kya, LGM) was based on the circulation model of CCSM4. The numbers in blue between pairs of modeled distribution indicate the Schoner’D values, where higher values represent higher niche similarity.

**Supplementary Fig. S7.**
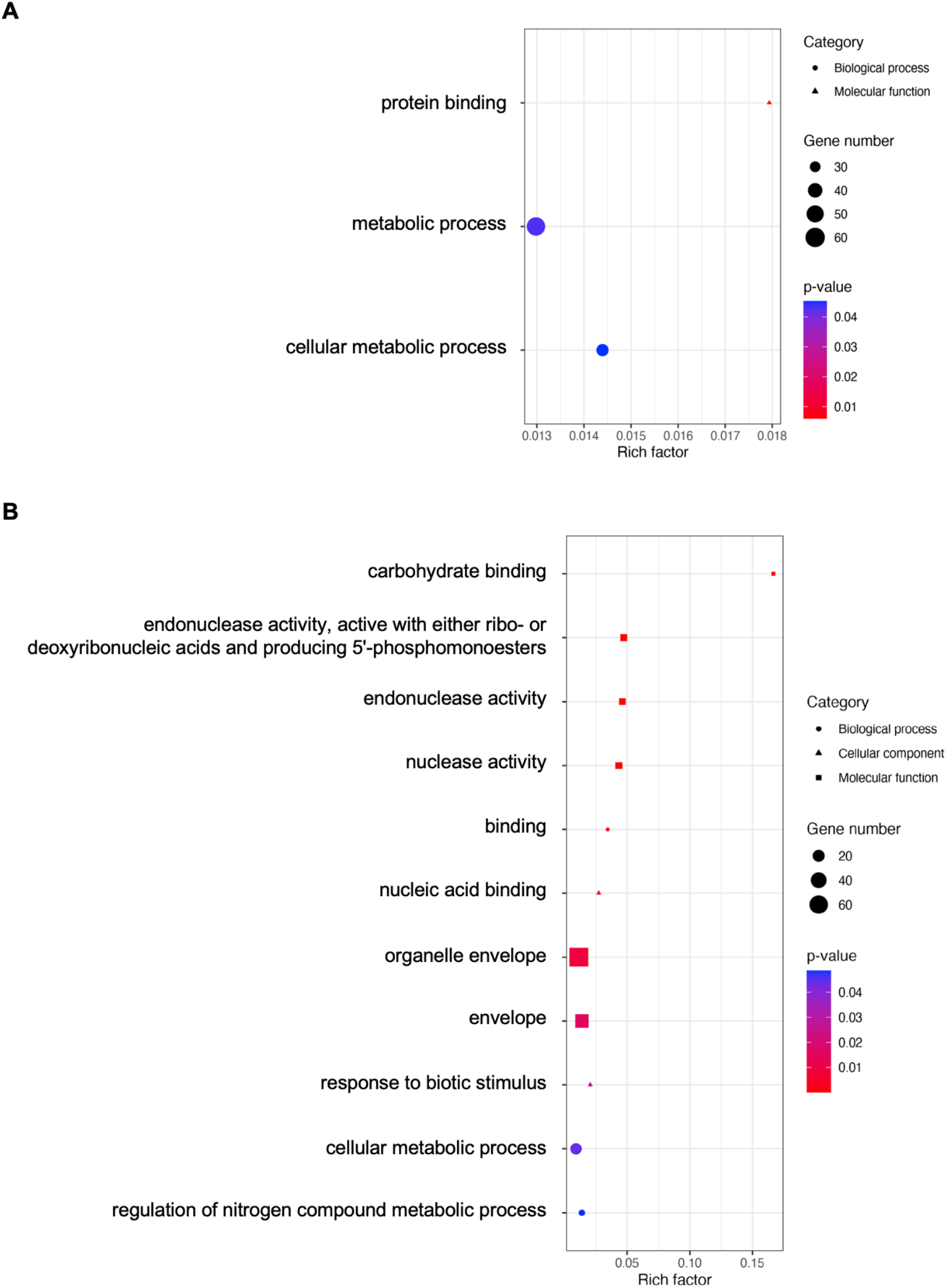
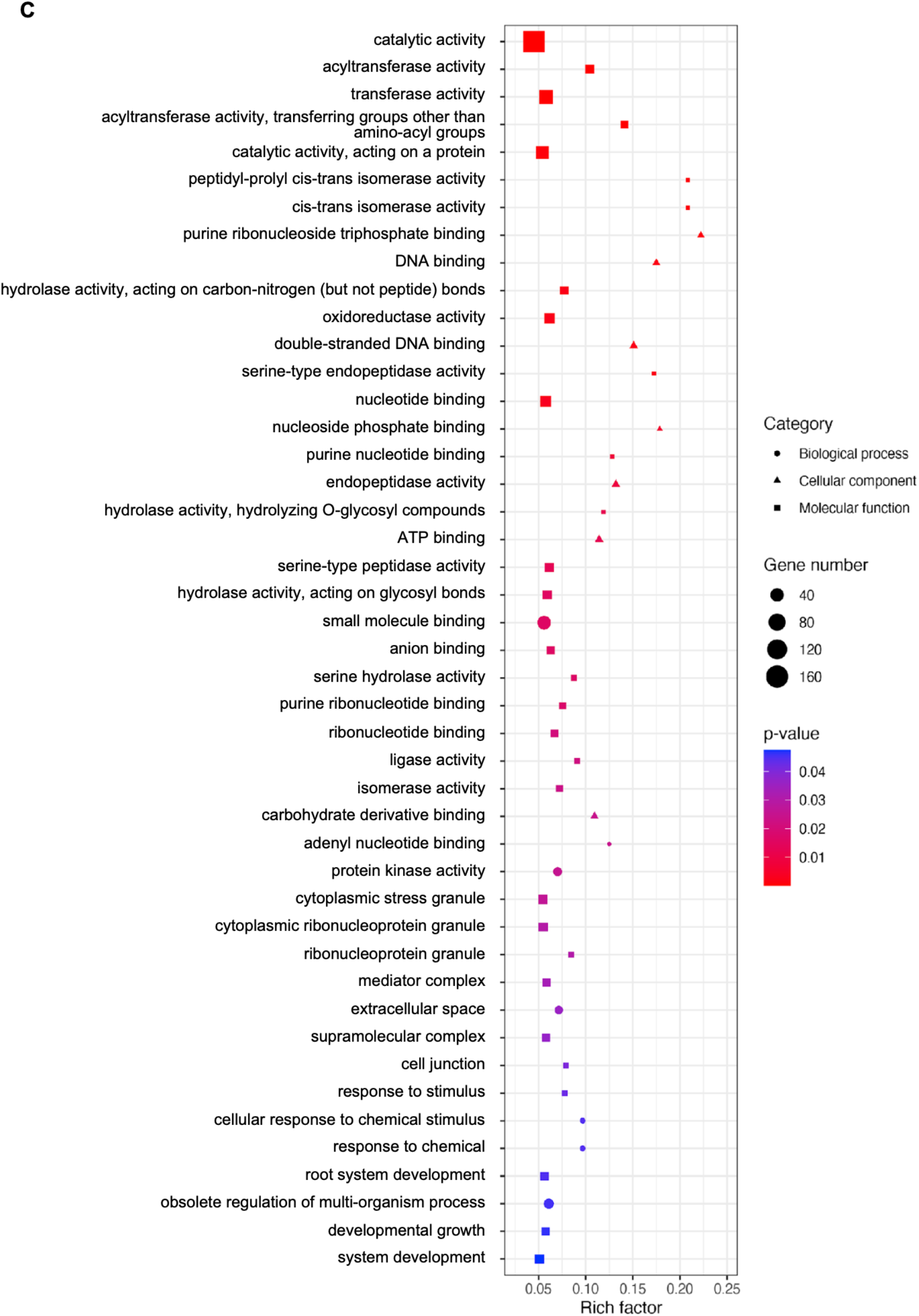
Gene Ontology (GO) enrichment of candidate genes in the regions of signatures of selection. Shown are the GO enrichment of candidate selective sweep genes from the (A) composite likelihood ratio (CLR) of the wild group, (B) CLR of the cultivars, and (C) reduction of diversity.

**Supplementary Fig. S8.**
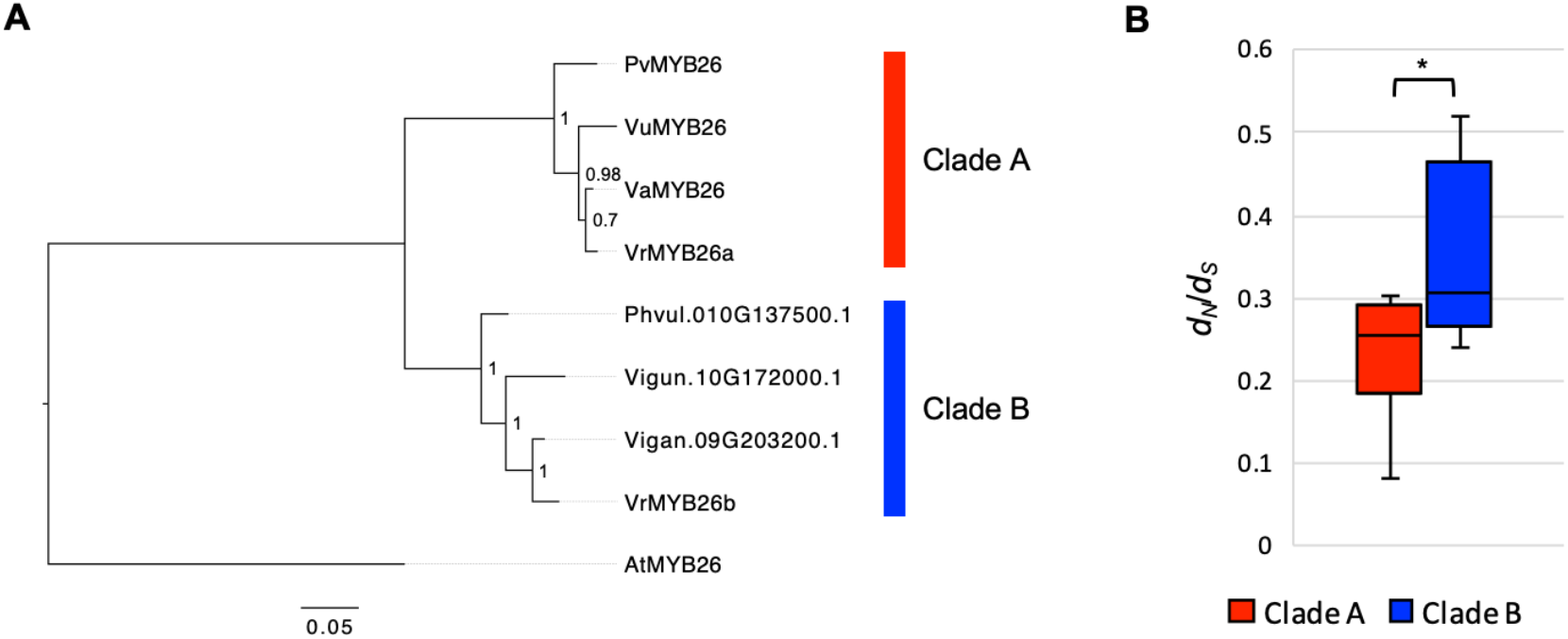
Evolution of *MYB26* homologs in legumes. (A) Maximum-likelihood (ML) tree constructed from the coding sequences of the *MYB26* homologs using *AtMYB26* as an outgroup. The numbers adjacent to nodes indicate the proportion of support in 1,000 bootstrap re-samplings. (B) The *d_N_*/*d_S_* ratios of the *MYB26* orthologs are estimated based on the pairwise comparison in each of the two clades in the ML tree. The boxes indicate medians and interquartile ranges; the whiskers indicate 95% values (* indicates *p* value < 0.05).

**Supplementary Fig. S9.**
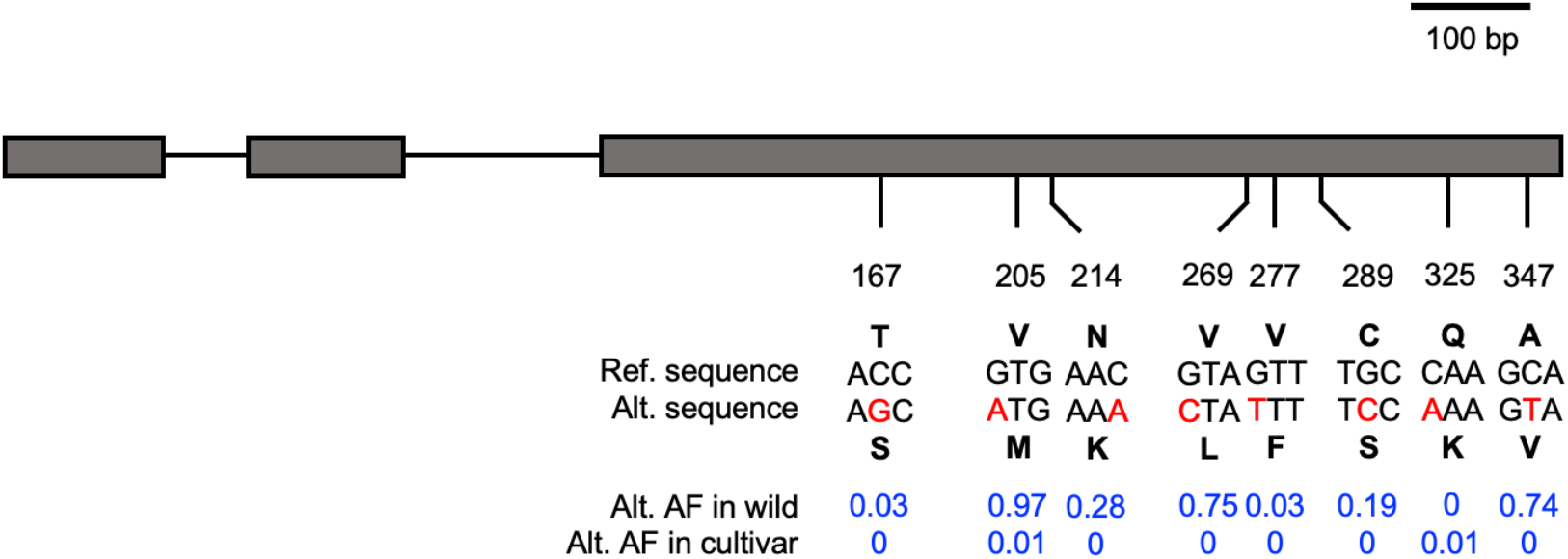
Diagram of the *VrMYB26a* non-synonymous changes. Shown are the positions of non-synonymous SNPs and the frequency of alternative alleles (Alt. AF) in wild and cultivated populations.

**Supplementary Fig. S10.**
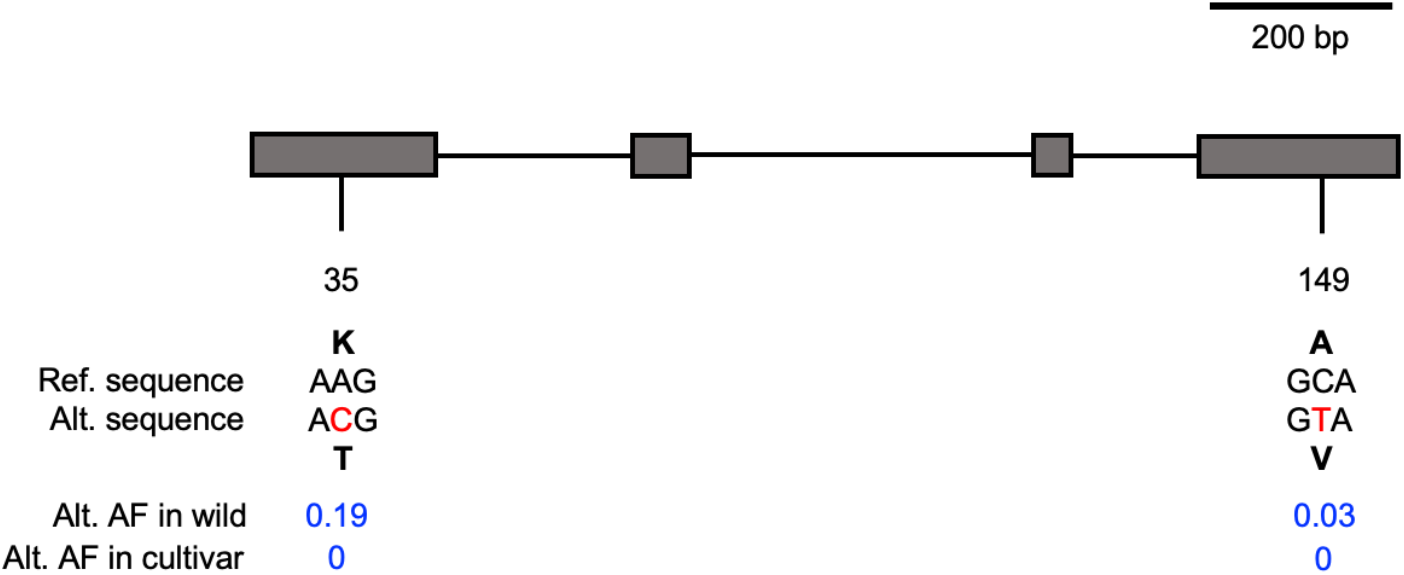
Diagram of the *VrDet1* non-synonymous changes. Shown are the positions of non-synonymous SNPs and the frequency of alternative alleles (Alt. AF) in wild and cultivated populations.

**Supplementary Fig. S11.**
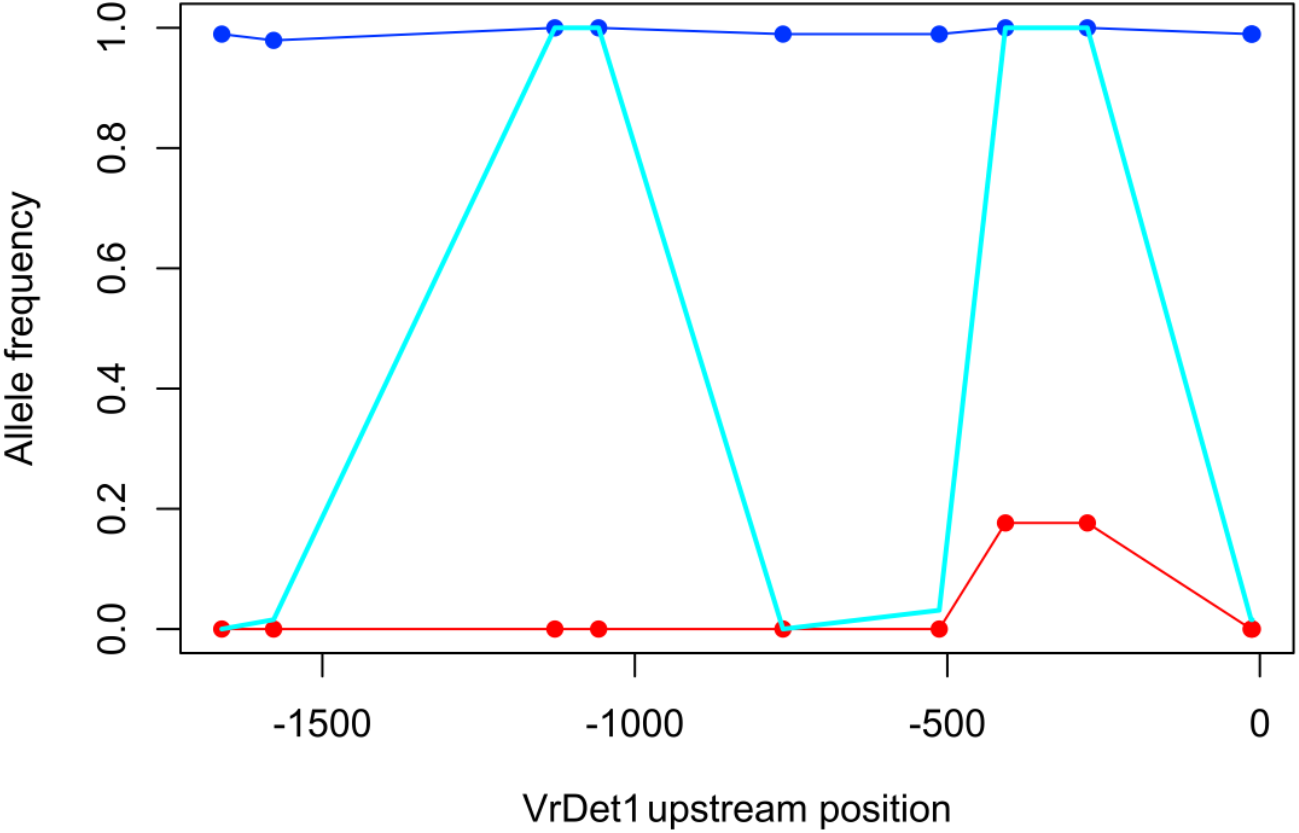
Distribution of allele frequency in *VrDet1* upstream 2kb region among three clades (red: Clade I, sky-blue: Clade II, blue: Clade III). Nine diagnostic SNPs with the large allele frequency differences between Clade I and III were shown, and the major allele in Clade III was used to estimate allele frequency.

**Supplementary Table S1.**
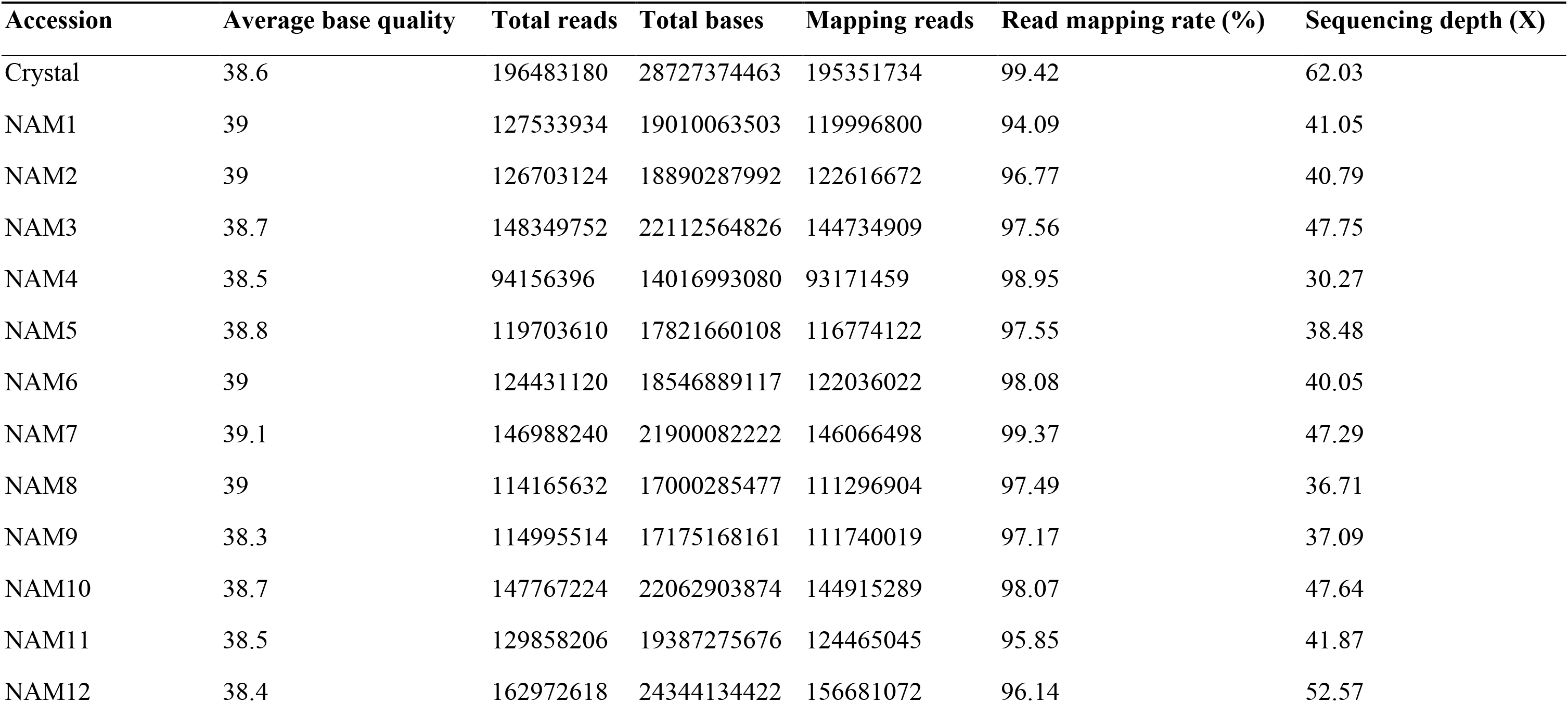
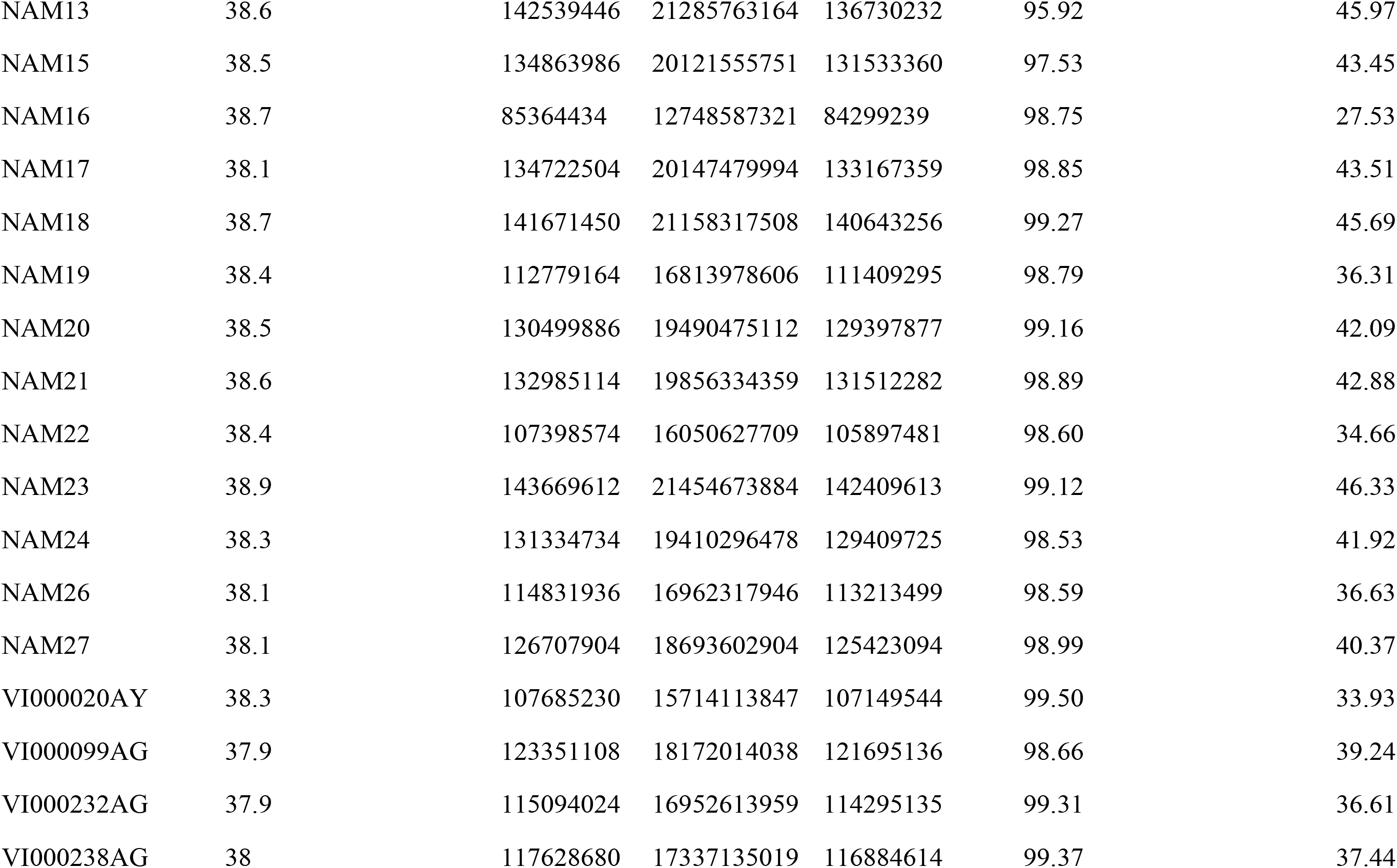
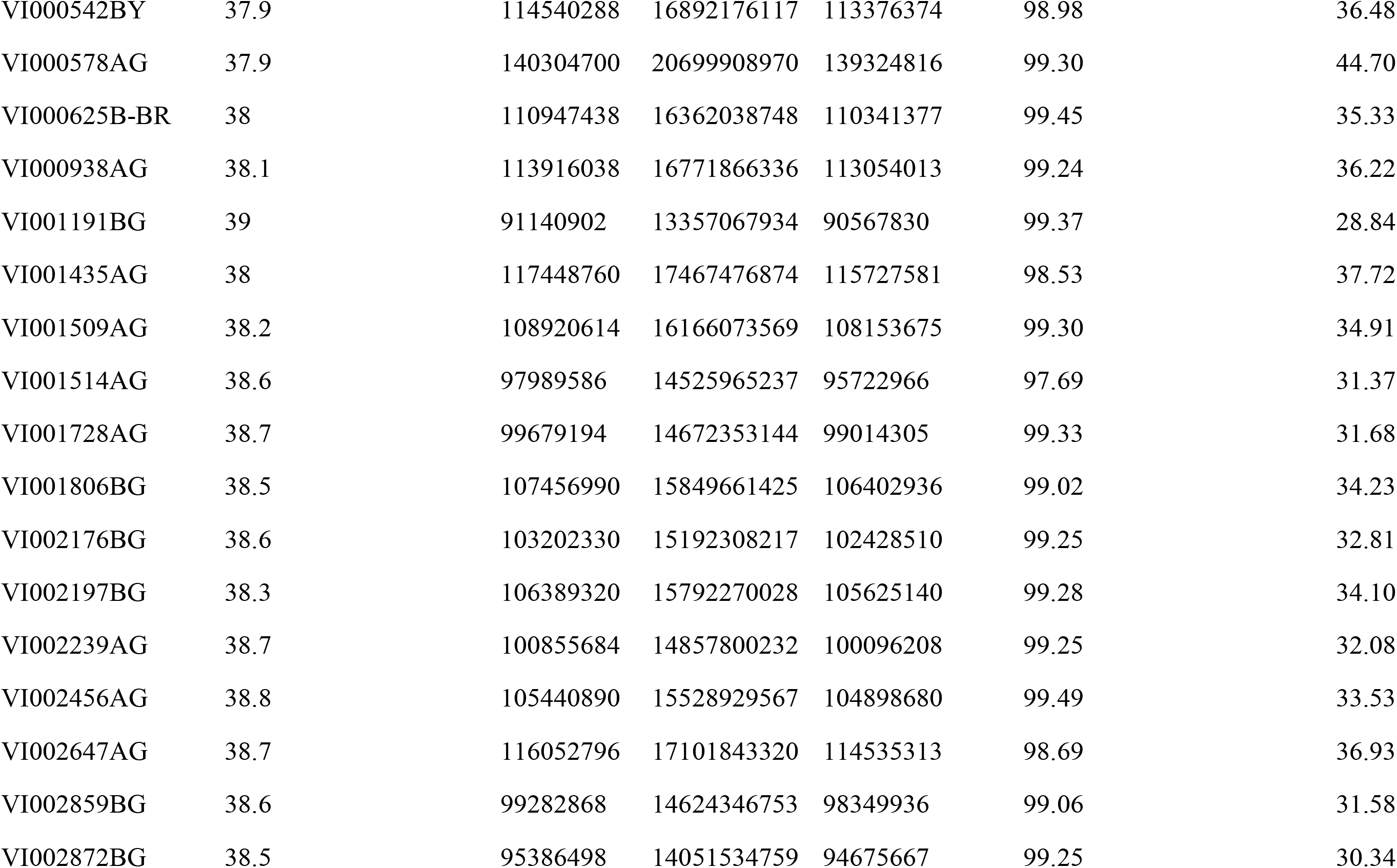
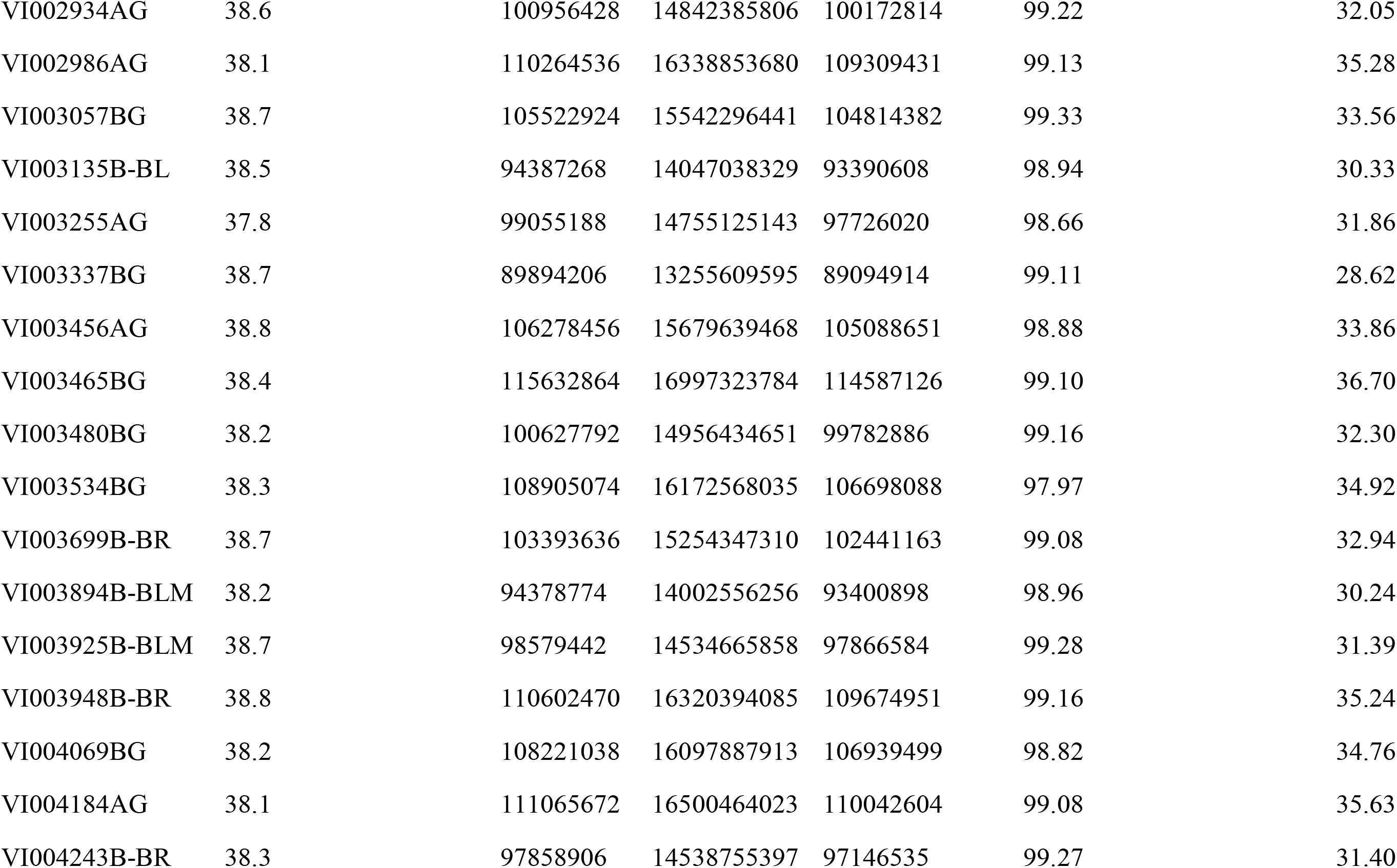
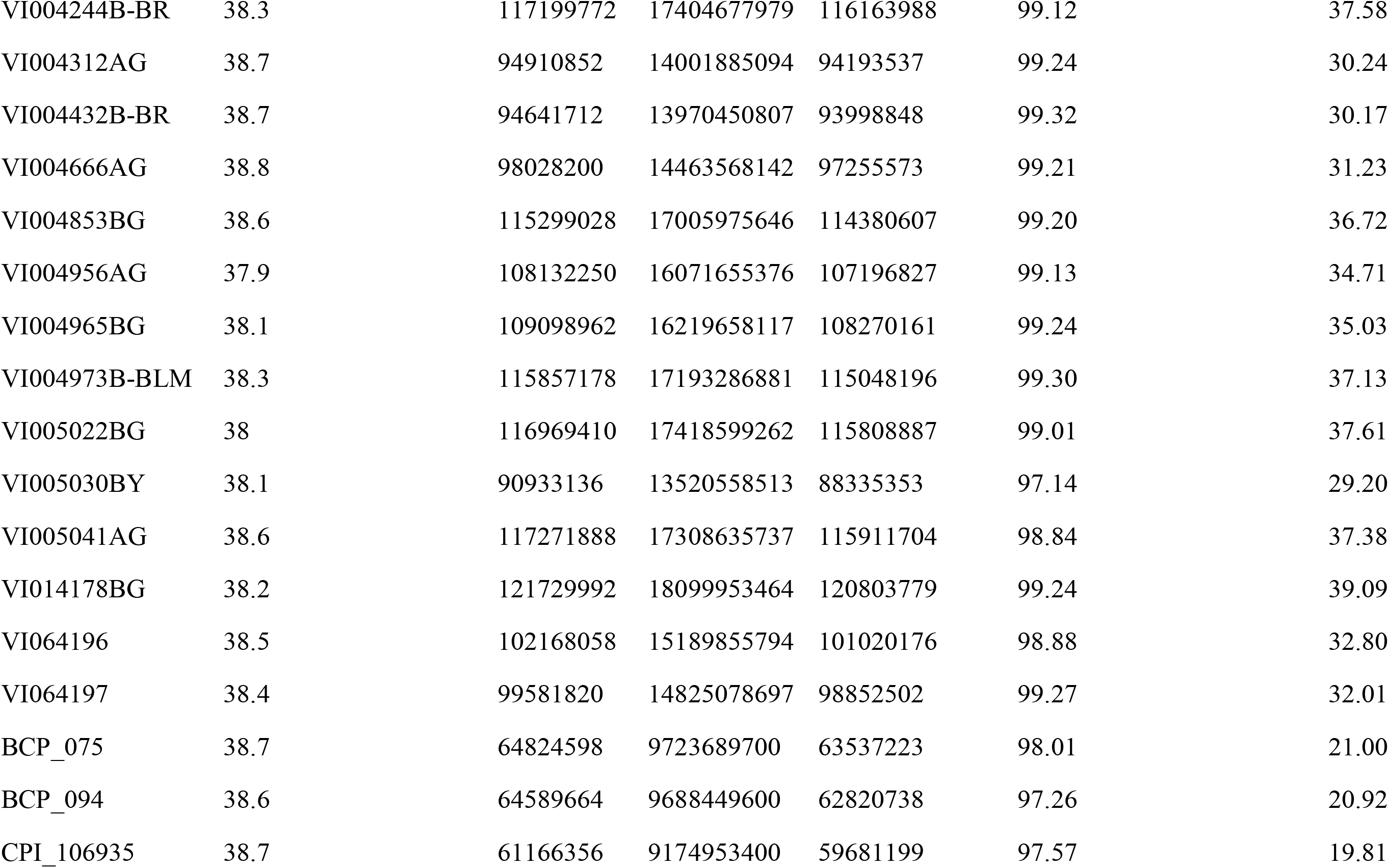
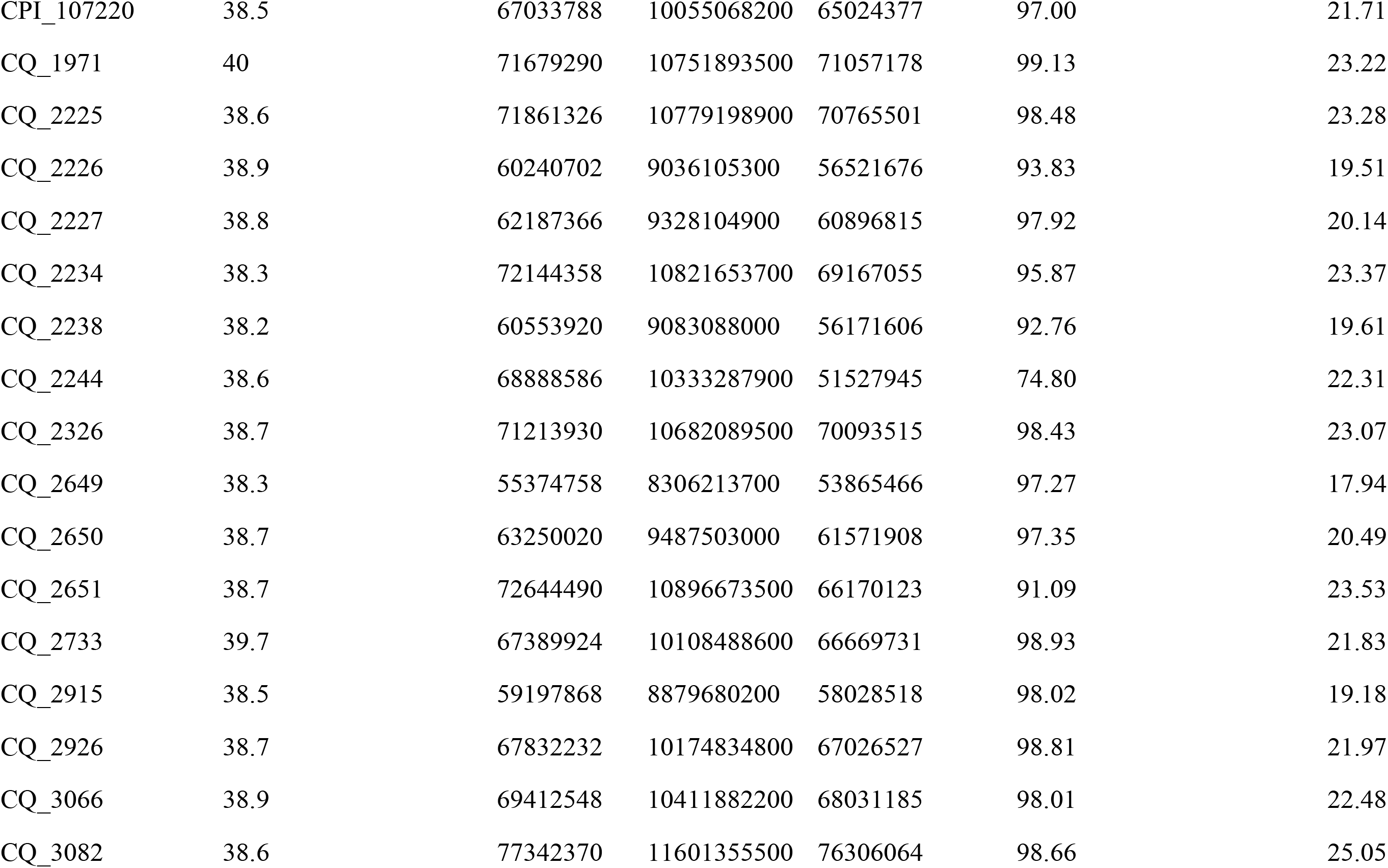
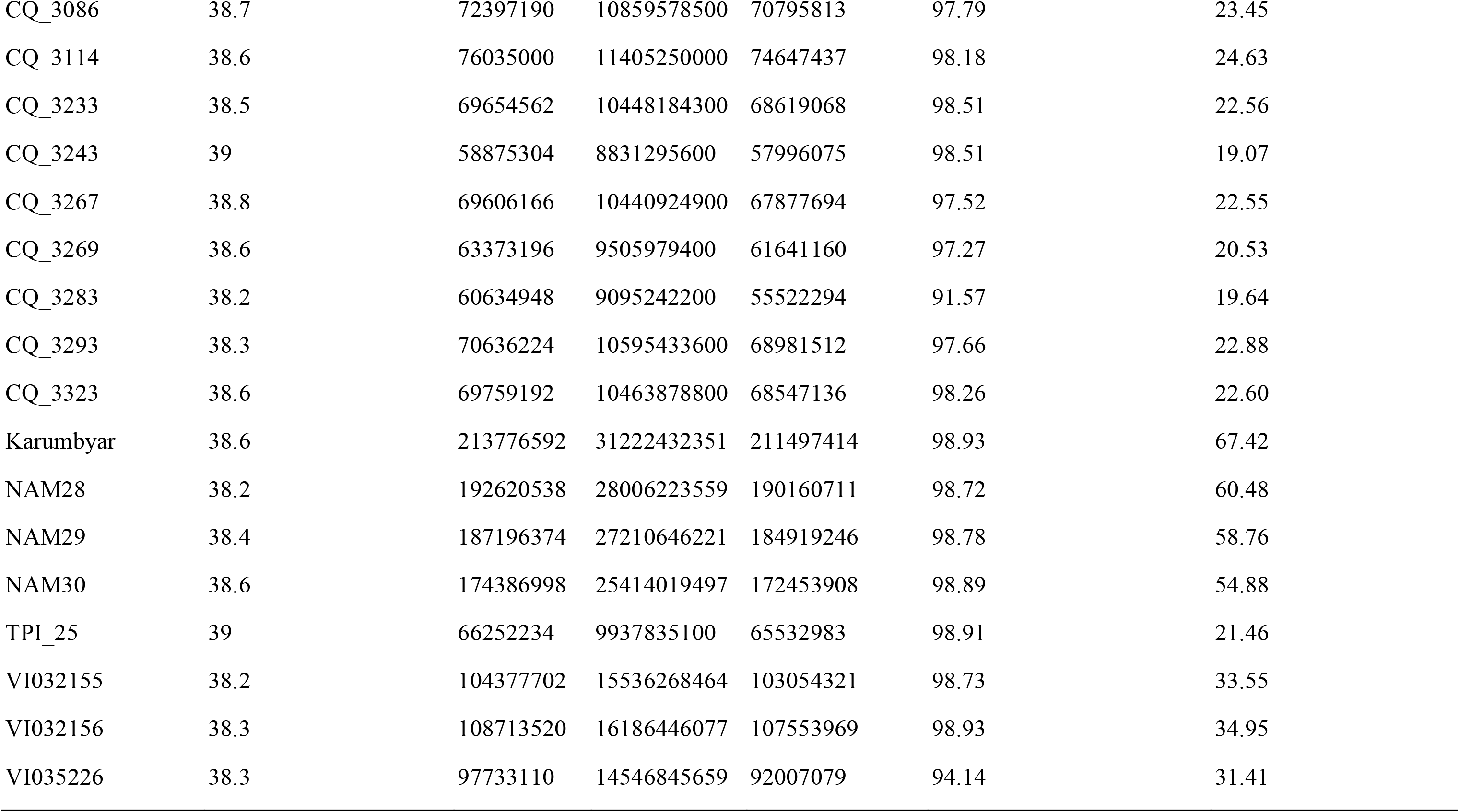
Summary of sequencing data for each accession analyzed in this study.

| Accession | Average base quality | Total reads | Total bases | Mapping reads | Read mapping rate (%) | Sequencing depth (X) |
| --- | --- | --- | --- | --- | --- | --- |
| Crystal | 38.6 | 196483180 | 28727374463 | 195351734 | 99.42 | 62.03 |
| NAM1 | 39 | 127533934 | 19010063503 | 119996800 | 94.09 | 41.05 |
| NAM2 | 39 | 126703124 | 18890287992 | 122616672 | 96.77 | 40.79 |
| NAM3 | 38.7 | 148349752 | 22112564826 | 144734909 | 97.56 | 47.75 |
| NAM4 | 38.5 | 94156396 | 14016993080 | 93171459 | 98.95 | 30.27 |
| NAM5 | 38.8 | 119703610 | 17821660108 | 116774122 | 97.55 | 38.48 |
| NAM6 | 39 | 124431120 | 18546889117 | 122036022 | 98.08 | 40.05 |
| NAM7 | 39.1 | 146988240 | 21900082222 | 146066498 | 99.37 | 47.29 |
| NAM8 | 39 | 114165632 | 17000285477 | 111296904 | 97.49 | 36.71 |
| NAM9 | 38.3 | 114995514 | 17175168161 | 111740019 | 97.17 | 37.09 |
| NAM10 | 38.7 | 147767224 | 22062903874 | 144915289 | 98.07 | 47.64 |
| NAM11 | 38.5 | 129858206 | 19387275676 | 124465045 | 95.85 | 41.87 |
| NAM12 | 38.4 | 162972618 | 24344134422 | 156681072 | 96.14 | 52.57 |
| NAM13 | 38.6 | 142539446 | 21285763164 | 136730232 | 95.92 | 45.97 |
| NAM15 | 38.5 | 134863986 | 20121555751 | 131533360 | 97.53 | 43.45 |
| NAM16 | 38.7 | 85364434 | 12748587321 | 84299239 | 98.75 | 27.53 |
| NAM17 | 38.1 | 134722504 | 20147479994 | 133167359 | 98.85 | 43.51 |
| NAM18 | 38.7 | 141671450 | 21158317508 | 140643256 | 99.27 | 45.69 |
| NAM19 | 38.4 | 112779164 | 16813978606 | 111409295 | 98.79 | 36.31 |
| NAM20 | 38.5 | 130499886 | 19490475112 | 129397877 | 99.16 | 42.09 |
| NAM21 | 38.6 | 132985114 | 19856334359 | 131512282 | 98.89 | 42.88 |
| NAM22 | 38.4 | 107398574 | 16050627709 | 105897481 | 98.60 | 34.66 |
| NAM23 | 38.9 | 143669612 | 21454673884 | 142409613 | 99.12 | 46.33 |
| NAM24 | 38.3 | 131334734 | 19410296478 | 129409725 | 98.53 | 41.92 |
| NAM26 | 38.1 | 114831936 | 16962317946 | 113213499 | 98.59 | 36.63 |
| NAM27 | 38.1 | 126707904 | 18693602904 | 125423094 | 98.99 | 40.37 |
| VI000020AY | 38.3 | 107685230 | 15714113847 | 107149544 | 99.50 | 33.93 |
| VI000099AG | 37.9 | 123351108 | 18172014038 | 121695136 | 98.66 | 39.24 |
| VI000232AG | 37.9 | 115094024 | 16952613959 | 114295135 | 99.31 | 36.61 |
| VI000238AG | 38 | 117628680 | 17337135019 | 116884614 | 99.37 | 37.44 |
| VI000542BY | 37.9 | 114540288 | 16892176117 | 113376374 | 98.98 | 36.48 |
| VI000578AG | 37.9 | 140304700 | 20699908970 | 139324816 | 99.30 | 44.70 |
| VI000625B-BR | 38 | 110947438 | 16362038748 | 110341377 | 99.45 | 35.33 |
| VI000938AG | 38.1 | 113916038 | 16771866336 | 113054013 | 99.24 | 36.22 |
| VI001191BG | 39 | 91140902 | 13357067934 | 90567830 | 99.37 | 28.84 |
| VI001435AG | 38 | 117448760 | 17467476874 | 115727581 | 98.53 | 37.72 |
| VI001509AG | 38.2 | 108920614 | 16166073569 | 108153675 | 99.30 | 34.91 |
| VI001514AG | 38.6 | 97989586 | 14525965237 | 95722966 | 97.69 | 31.37 |
| VI001728AG | 38.7 | 99679194 | 14672353144 | 99014305 | 99.33 | 31.68 |
| VI001806BG | 38.5 | 107456990 | 15849661425 | 106402936 | 99.02 | 34.23 |
| VI002176BG | 38.6 | 103202330 | 15192308217 | 102428510 | 99.25 | 32.81 |
| VI002197BG | 38.3 | 106389320 | 15792270028 | 105625140 | 99.28 | 34.10 |
| VI002239AG | 38.7 | 100855684 | 14857800232 | 100096208 | 99.25 | 32.08 |
| VI002456AG | 38.8 | 105440890 | 15528929567 | 104898680 | 99.49 | 33.53 |
| VI002647AG | 38.7 | 116052796 | 17101843320 | 114535313 | 98.69 | 36.93 |
| VI002859BG | 38.6 | 99282868 | 14624346753 | 98349936 | 99.06 | 31.58 |
| VI002872BG | 38.5 | 95386498 | 14051534759 | 94675667 | 99.25 | 30.34 |
| VI002934AG | 38.6 | 100956428 | 14842385806 | 100172814 | 99.22 | 32.05 |
| VI002986AG | 38.1 | 110264536 | 16338853680 | 109309431 | 99.13 | 35.28 |
| VI003057BG | 38.7 | 105522924 | 15542296441 | 104814382 | 99.33 | 33.56 |
| VI003135B-BL | 38.5 | 94387268 | 14047038329 | 93390608 | 98.94 | 30.33 |
| VI003255AG | 37.8 | 99055188 | 14755125143 | 97726020 | 98.66 | 31.86 |
| VI003337BG | 38.7 | 89894206 | 13255609595 | 89094914 | 99.11 | 28.62 |
| VI003456AG | 38.8 | 106278456 | 15679639468 | 105088651 | 98.88 | 33.86 |
| VI003465BG | 38.4 | 115632864 | 16997323784 | 114587126 | 99.10 | 36.70 |
| VI003480BG | 38.2 | 100627792 | 14956434651 | 99782886 | 99.16 | 32.30 |
| VI003534BG | 38.3 | 108905074 | 16172568035 | 106698088 | 97.97 | 34.92 |
| VI003699B-BR | 38.7 | 103393636 | 15254347310 | 102441163 | 99.08 | 32.94 |
| VI003894B-BLM | 38.2 | 94378774 | 14002556256 | 93400898 | 98.96 | 30.24 |
| VI003925B-BLM | 38.7 | 98579442 | 14534665858 | 97866584 | 99.28 | 31.39 |
| VI003948B-BR | 38.8 | 110602470 | 16320394085 | 109674951 | 99.16 | 35.24 |
| VI004069BG | 38.2 | 108221038 | 16097887913 | 106939499 | 98.82 | 34.76 |
| VI004184AG | 38.1 | 111065672 | 16500464023 | 110042604 | 99.08 | 35.63 |
| VI004243B-BR | 38.3 | 97858906 | 14538755397 | 97146535 | 99.27 | 31.40 |
| VI004244B-BR | 38.3 | 117199772 | 17404677979 | 116163988 | 99.12 | 37.58 |
| VI004312AG | 38.7 | 94910852 | 14001885094 | 94193537 | 99.24 | 30.24 |
| VI004432B-BR | 38.7 | 94641712 | 13970450807 | 93998848 | 99.32 | 30.17 |
| VI004666AG | 38.8 | 98028200 | 14463568142 | 97255573 | 99.21 | 31.23 |
| VI004853BG | 38.6 | 115299028 | 17005975646 | 114380607 | 99.20 | 36.72 |
| VI004956AG | 37.9 | 108132250 | 16071655376 | 107196827 | 99.13 | 34.71 |
| VI004965BG | 38.1 | 109098962 | 16219658117 | 108270161 | 99.24 | 35.03 |
| VI004973B-BLM | 38.3 | 115857178 | 17193286881 | 115048196 | 99.30 | 37.13 |
| VI005022BG | 38 | 116969410 | 17418599262 | 115808887 | 99.01 | 37.61 |
| VI005030BY | 38.1 | 90933136 | 13520558513 | 88335353 | 97.14 | 29.20 |
| VI005041AG | 38.6 | 117271888 | 17308635737 | 115911704 | 98.84 | 37.38 |
| VI014178BG | 38.2 | 121729992 | 18099953464 | 120803779 | 99.24 | 39.09 |
| VI064196 | 38.5 | 102168058 | 15189855794 | 101020176 | 98.88 | 32.80 |
| VI064197 | 38.4 | 99581820 | 14825078697 | 98852502 | 99.27 | 32.01 |
| BCP_075 | 38.7 | 64824598 | 9723689700 | 63537223 | 98.01 | 21.00 |
| BCP_094 | 38.6 | 64589664 | 9688449600 | 62820738 | 97.26 | 20.92 |
| CPI_106935 | 38.7 | 61166356 | 9174953400 | 59681199 | 97.57 | 19.81 |
| CPI_107220 | 38.5 | 67033788 | 10055068200 | 65024377 | 97.00 | 21.71 |
| CQ_1971 | 40 | 71679290 | 10751893500 | 71057178 | 99.13 | 23.22 |
| CQ_2225 | 38.6 | 71861326 | 10779198900 | 70765501 | 98.48 | 23.28 |
| CQ_2226 | 38.9 | 60240702 | 9036105300 | 56521676 | 93.83 | 19.51 |
| CQ_2227 | 38.8 | 62187366 | 9328104900 | 60896815 | 97.92 | 20.14 |
| CQ_2234 | 38.3 | 72144358 | 10821653700 | 69167055 | 95.87 | 23.37 |
| CQ_2238 | 38.2 | 60553920 | 9083088000 | 56171606 | 92.76 | 19.61 |
| CQ_2244 | 38.6 | 68888586 | 10333287900 | 51527945 | 74.80 | 22.31 |
| CQ_2326 | 38.7 | 71213930 | 10682089500 | 70093515 | 98.43 | 23.07 |
| CQ_2649 | 38.3 | 55374758 | 8306213700 | 53865466 | 97.27 | 17.94 |
| CQ_2650 | 38.7 | 63250020 | 9487503000 | 61571908 | 97.35 | 20.49 |
| CQ_2651 | 38.7 | 72644490 | 10896673500 | 66170123 | 91.09 | 23.53 |
| CQ_2733 | 39.7 | 67389924 | 10108488600 | 66669731 | 98.93 | 21.83 |
| CQ_2915 | 38.5 | 59197868 | 8879680200 | 58028518 | 98.02 | 19.18 |
| CQ_2926 | 38.7 | 67832232 | 10174834800 | 67026527 | 98.81 | 21.97 |
| CQ_3066 | 38.9 | 69412548 | 10411882200 | 68031185 | 98.01 | 22.48 |
| CQ_3082 | 38.6 | 77342370 | 11601355500 | 76306064 | 98.66 | 25.05 |
| CQ_3086 | 38.7 | 72397190 | 10859578500 | 70795813 | 97.79 | 23.45 |
| CQ_3114 | 38.6 | 76035000 | 11405250000 | 74647437 | 98.18 | 24.63 |
| CQ_3233 | 38.5 | 69654562 | 10448184300 | 68619068 | 98.51 | 22.56 |
| CQ_3243 | 39 | 58875304 | 8831295600 | 57996075 | 98.51 | 19.07 |
| CQ_3267 | 38.8 | 69606166 | 10440924900 | 67877694 | 97.52 | 22.55 |
| CQ_3269 | 38.6 | 63373196 | 9505979400 | 61641160 | 97.27 | 20.53 |
| CQ_3283 | 38.2 | 60634948 | 9095242200 | 55522294 | 91.57 | 19.64 |
| CQ_3293 | 38.3 | 70636224 | 10595433600 | 68981512 | 97.66 | 22.88 |
| CQ_3323 | 38.6 | 69759192 | 10463878800 | 68547136 | 98.26 | 22.60 |
| Karumbyar | 38.6 | 213776592 | 31222432351 | 211497414 | 98.93 | 67.42 |
| NAM28 | 38.2 | 192620538 | 28006223559 | 190160711 | 98.72 | 60.48 |
| NAM29 | 38.4 | 187196374 | 27210646221 | 184919246 | 98.78 | 58.76 |
| NAM30 | 38.6 | 174386998 | 25414019497 | 172453908 | 98.89 | 54.88 |
| TPI_25 | 39 | 66252234 | 9937835100 | 65532983 | 98.91 | 21.46 |
| VI032155 | 38.2 | 104377702 | 15536268464 | 103054321 | 98.73 | 33.55 |
| VI032156 | 38.3 | 108713520 | 16186446077 | 107553969 | 98.93 | 34.95 |
| VI035226 | 38.3 | 97733110 | 14546845659 | 92007079 | 94.14 | 31.41 |

**Supplementary Table S2.**
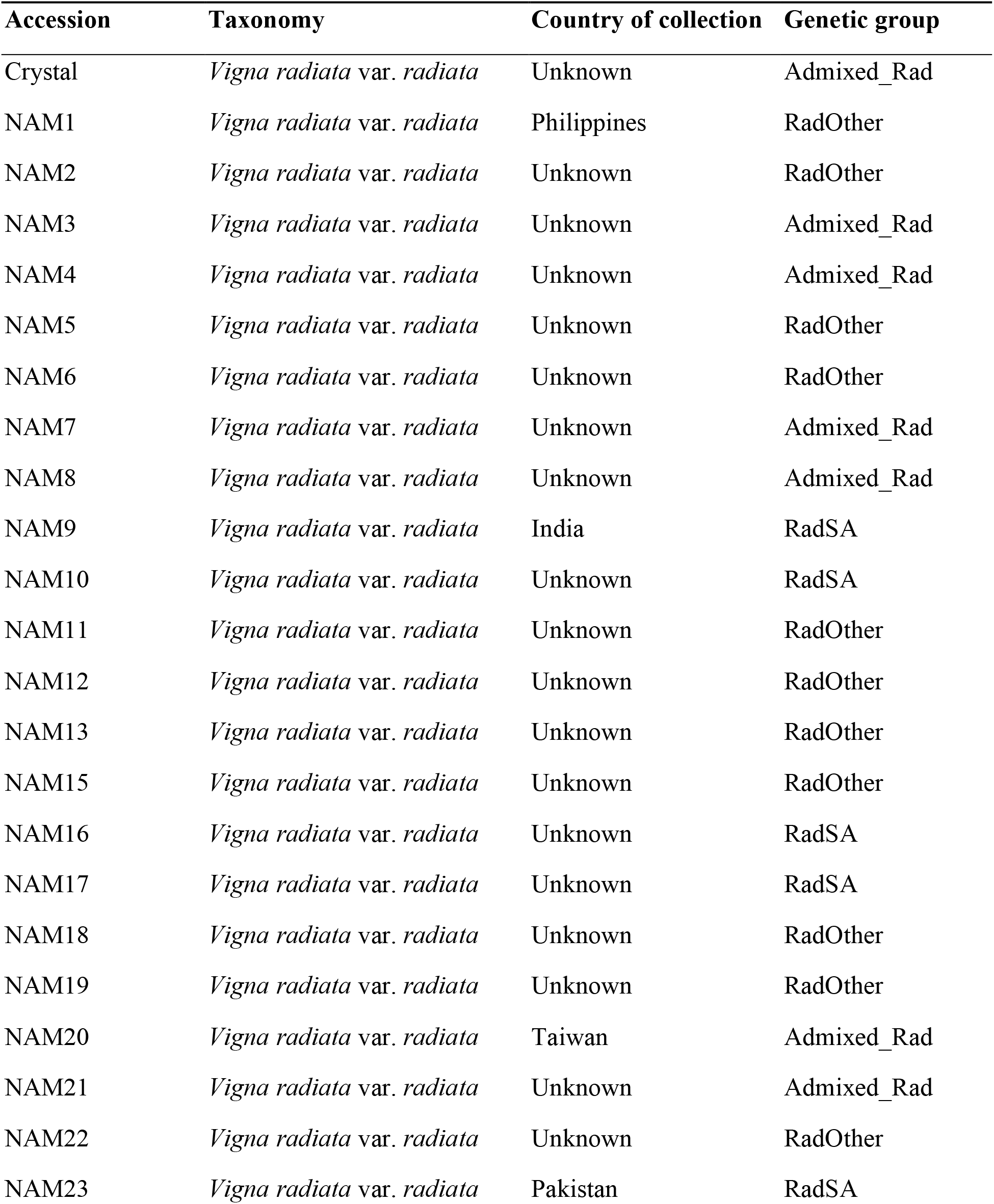
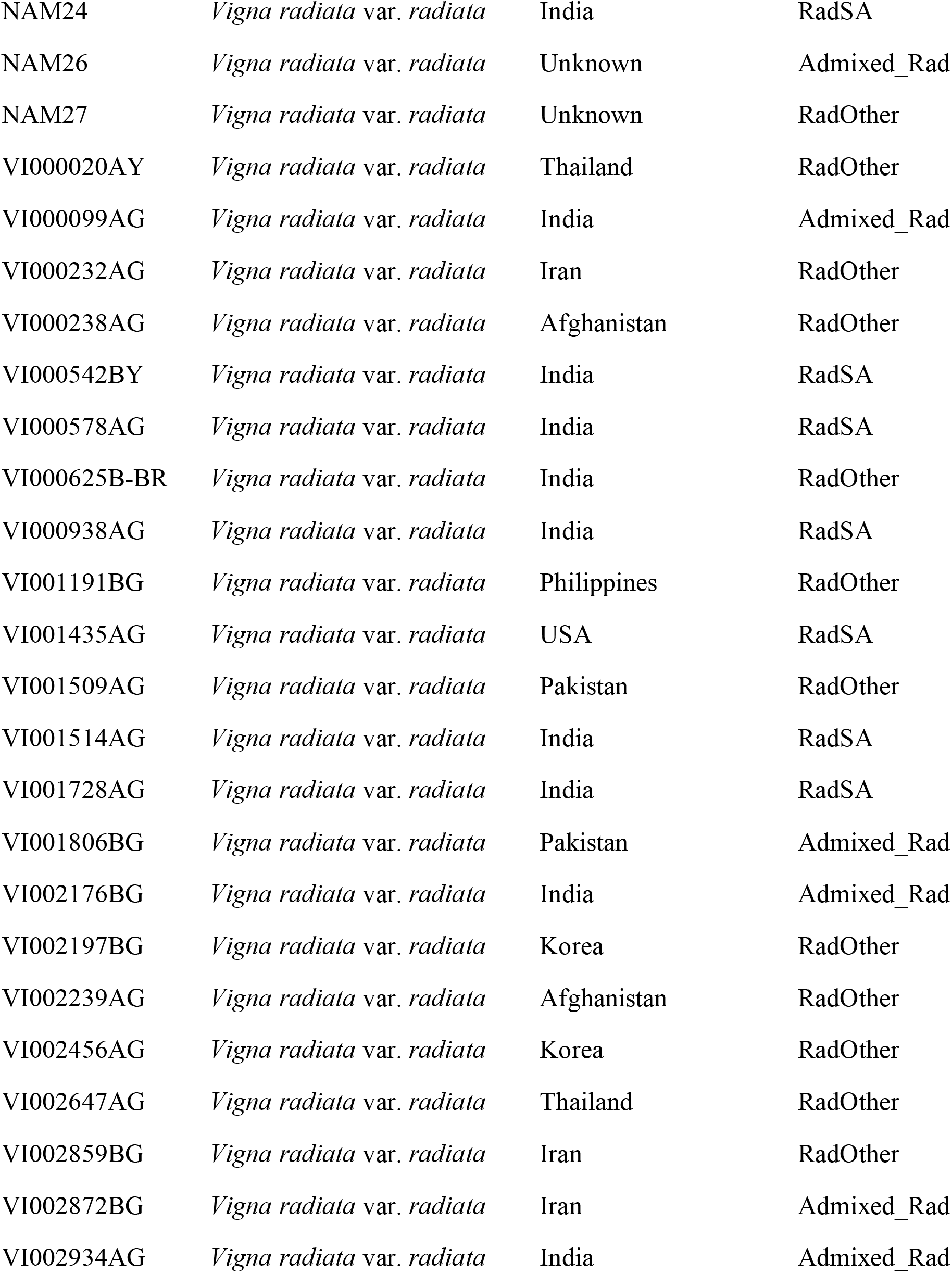
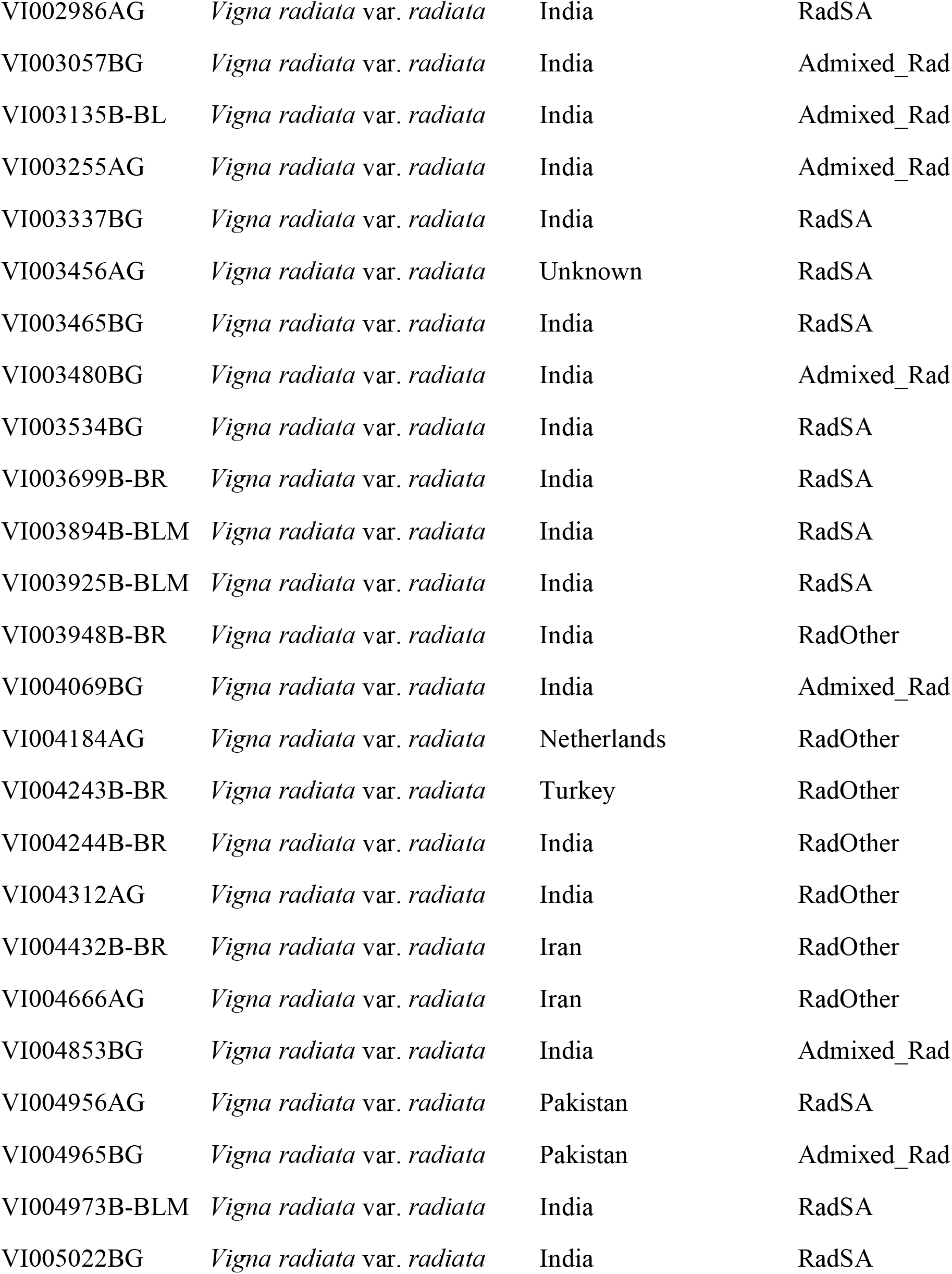
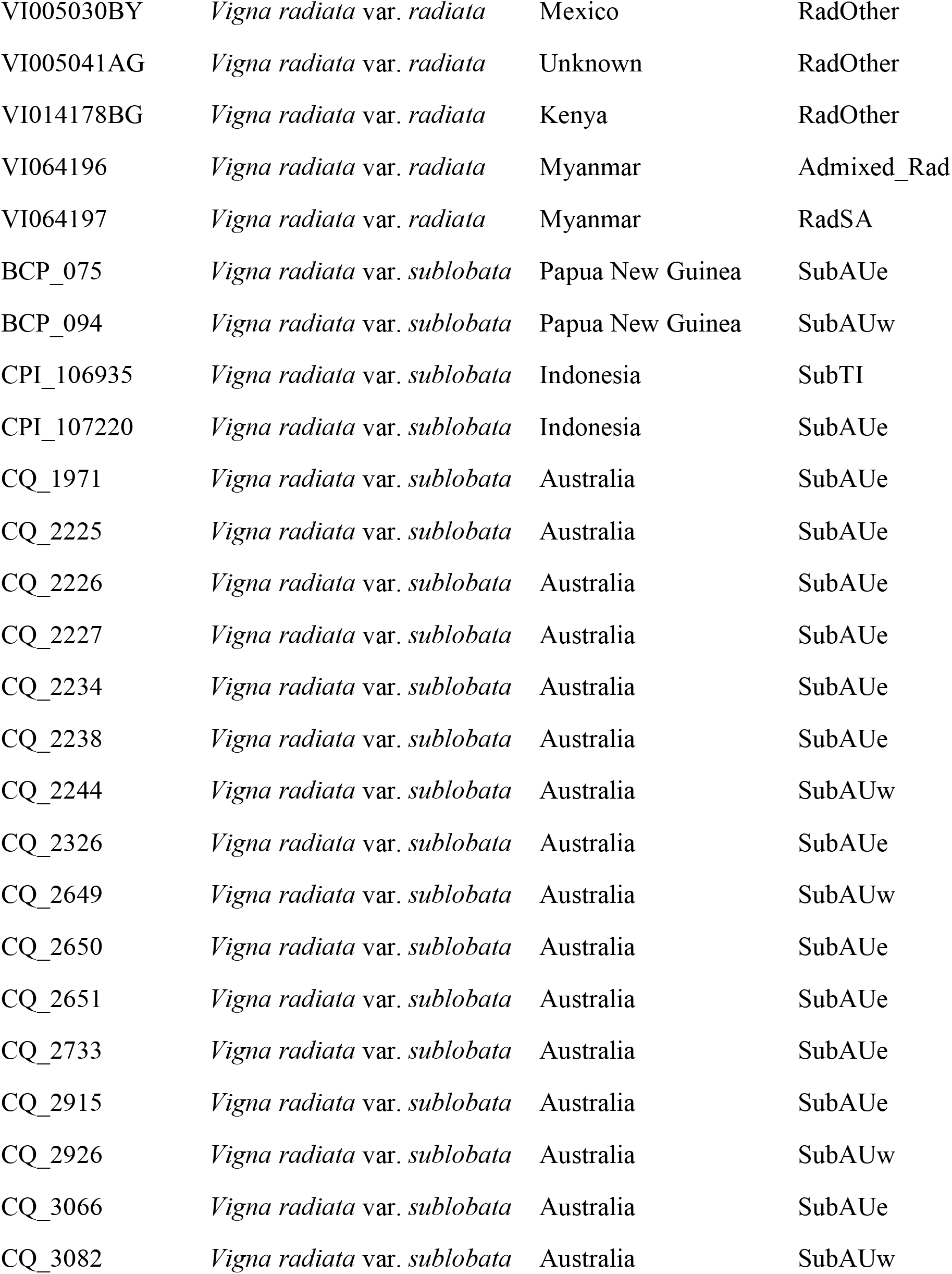
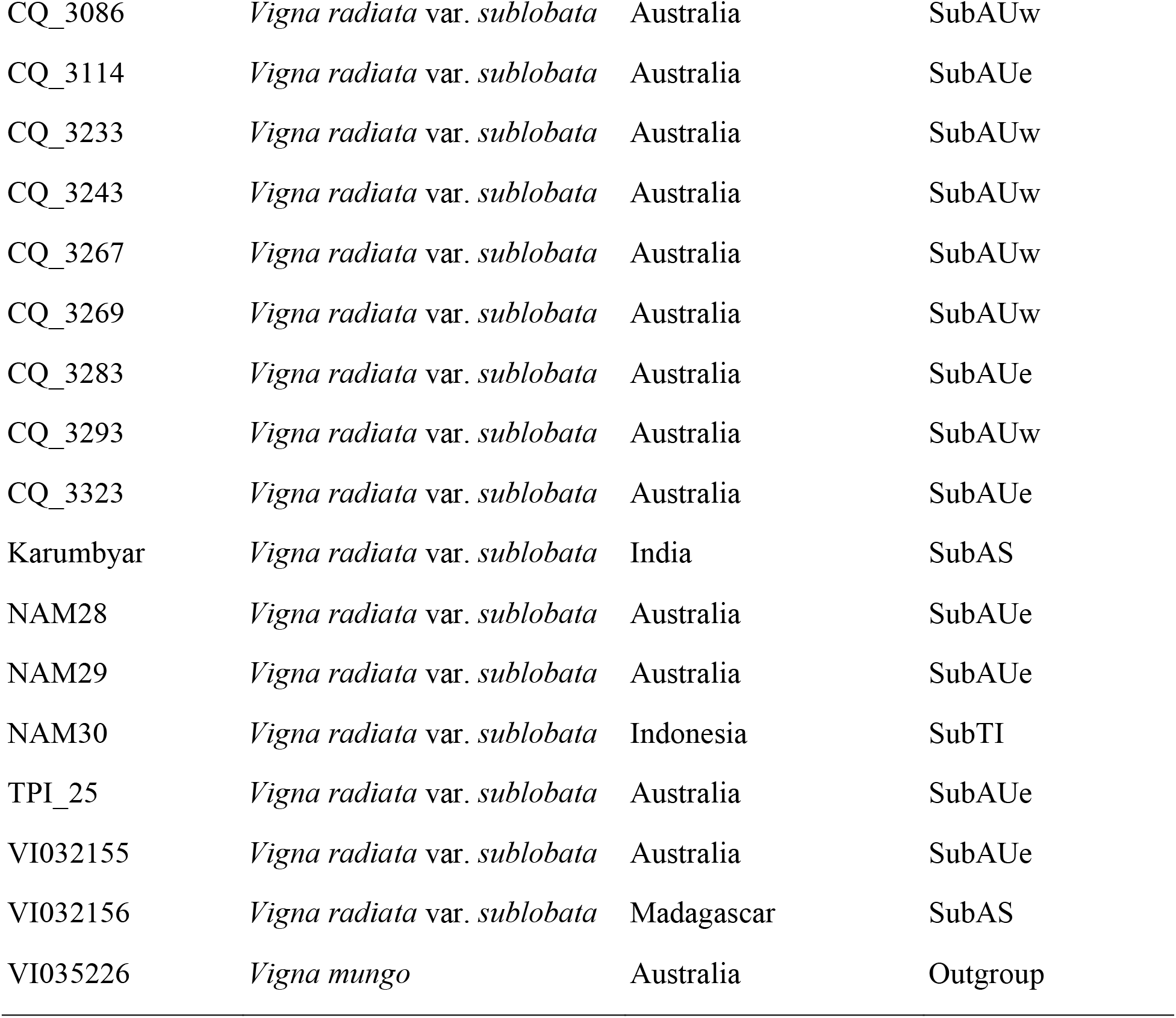
List of accessions used in this study.

**Supplementary Table S3.**
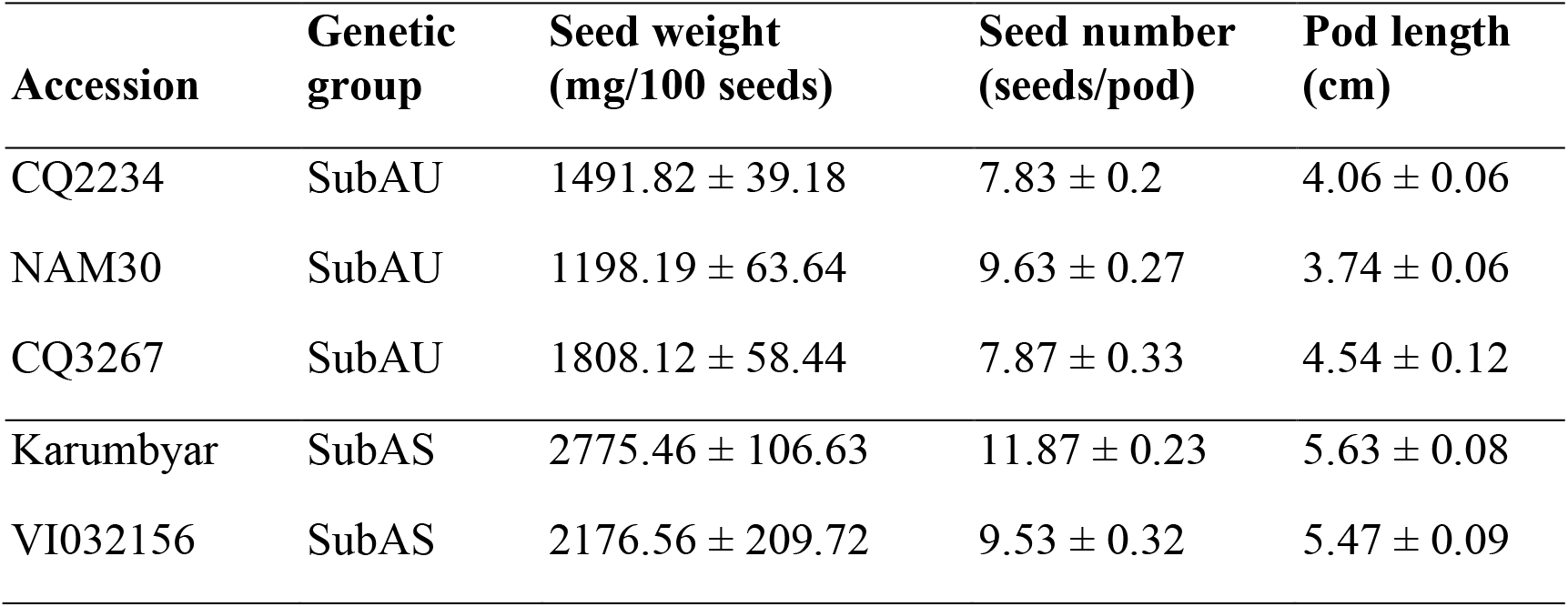
Measurement of seed weight in wild accessions from genetic groups of SubAU and SubAS. The data are presented as mean ± standard error.

**Supplementary Table S4.**
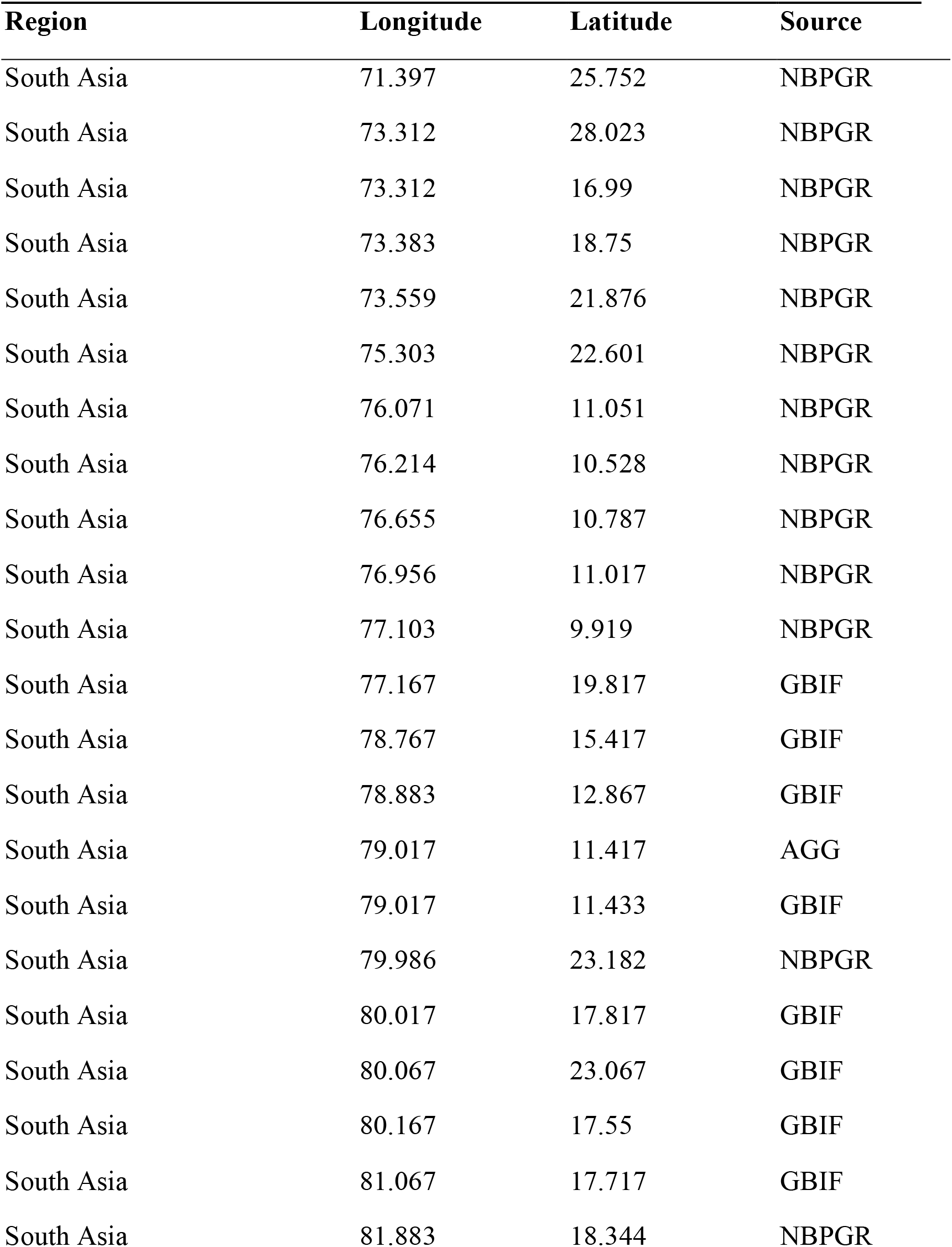
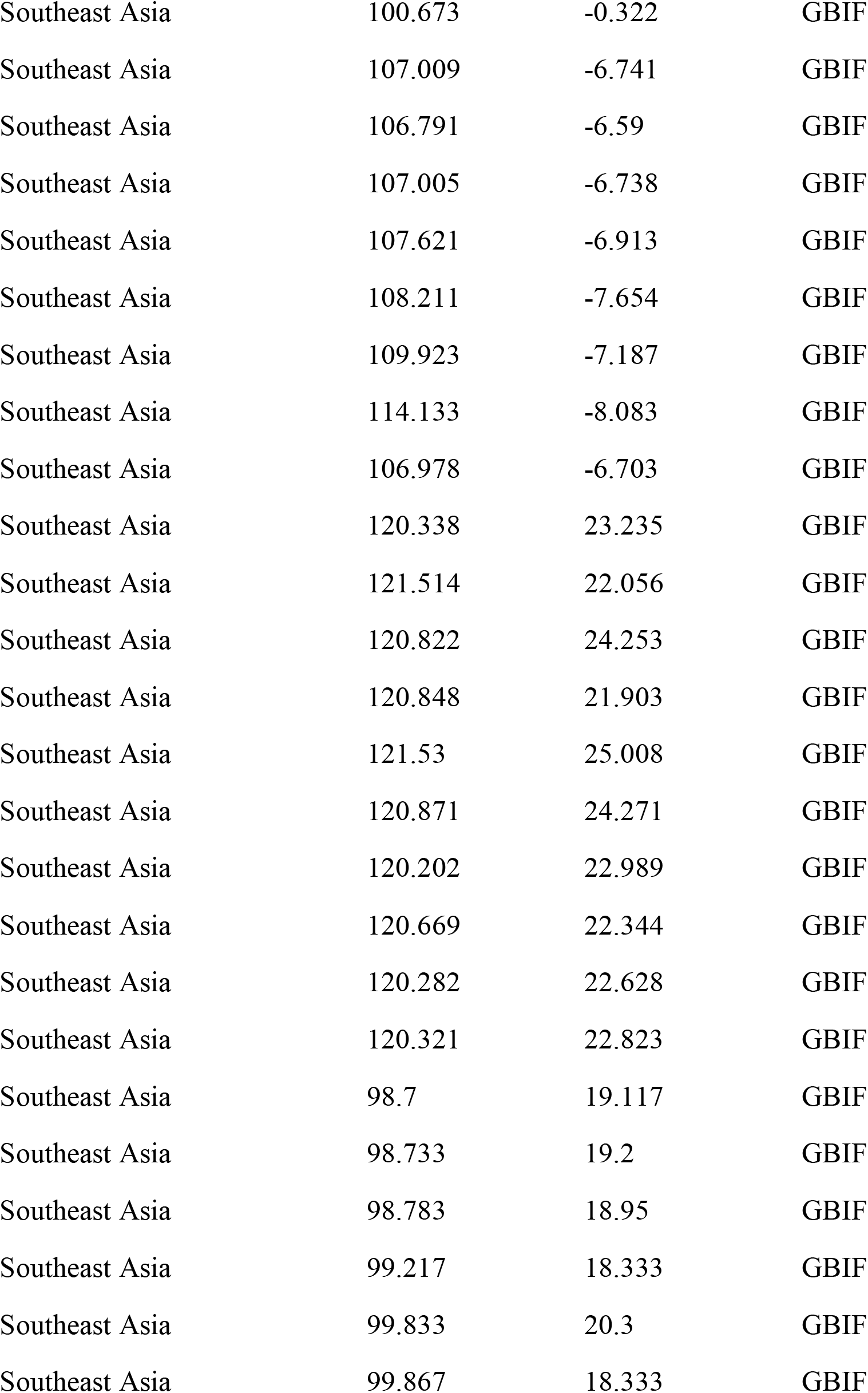
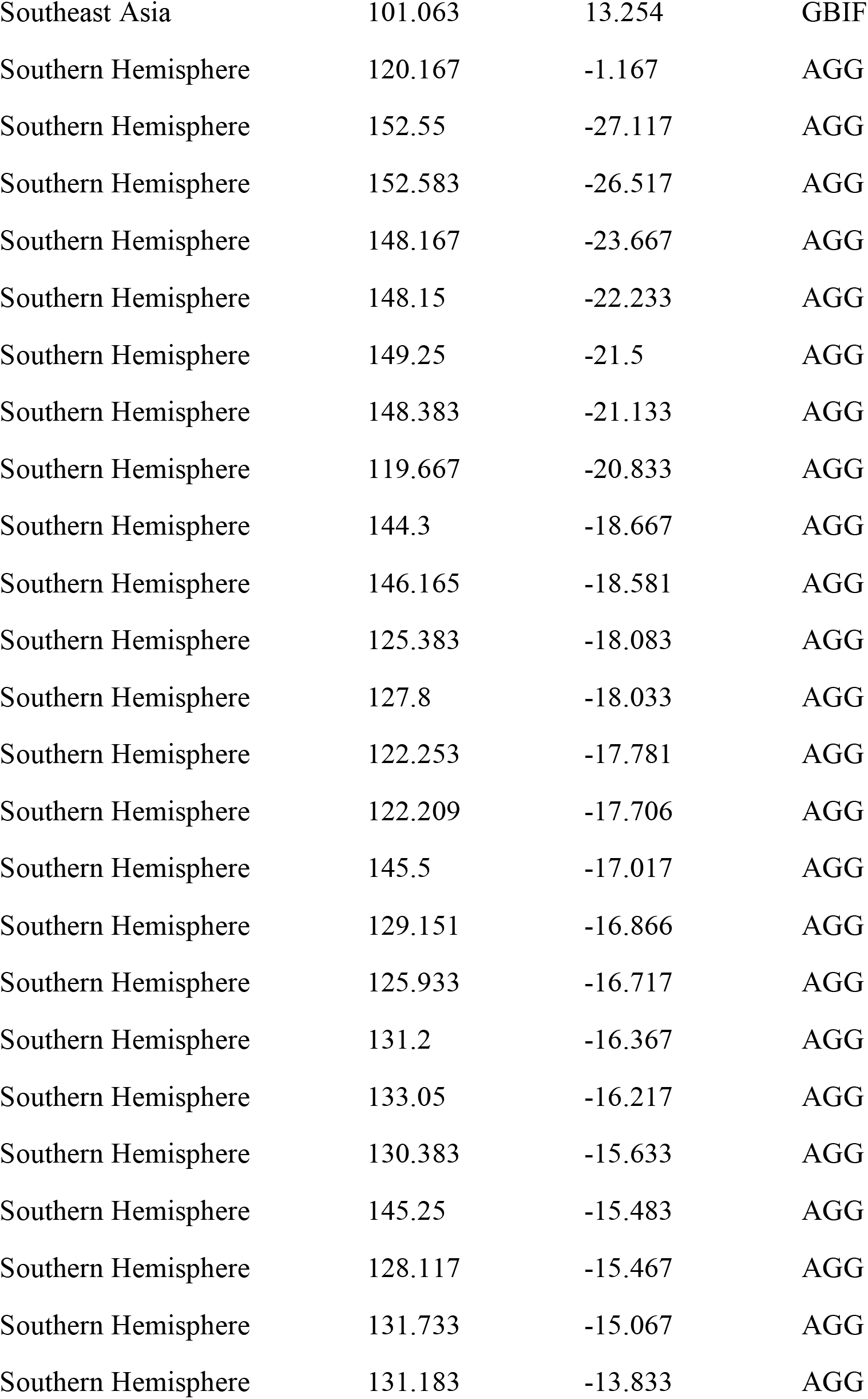
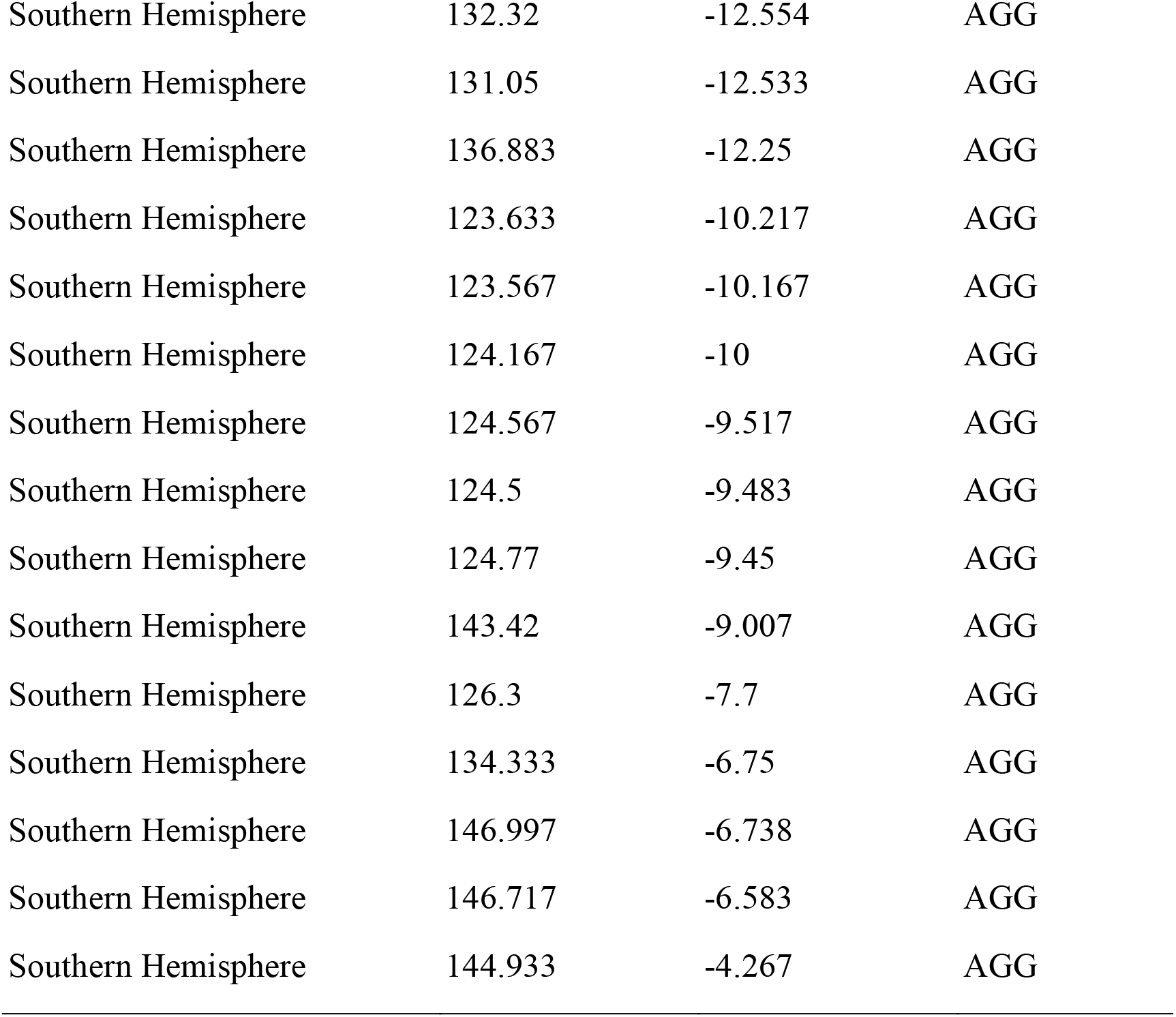
List of GPS sites of wild mungbeans used for niche modeling.

**Supplementary Table S5.**
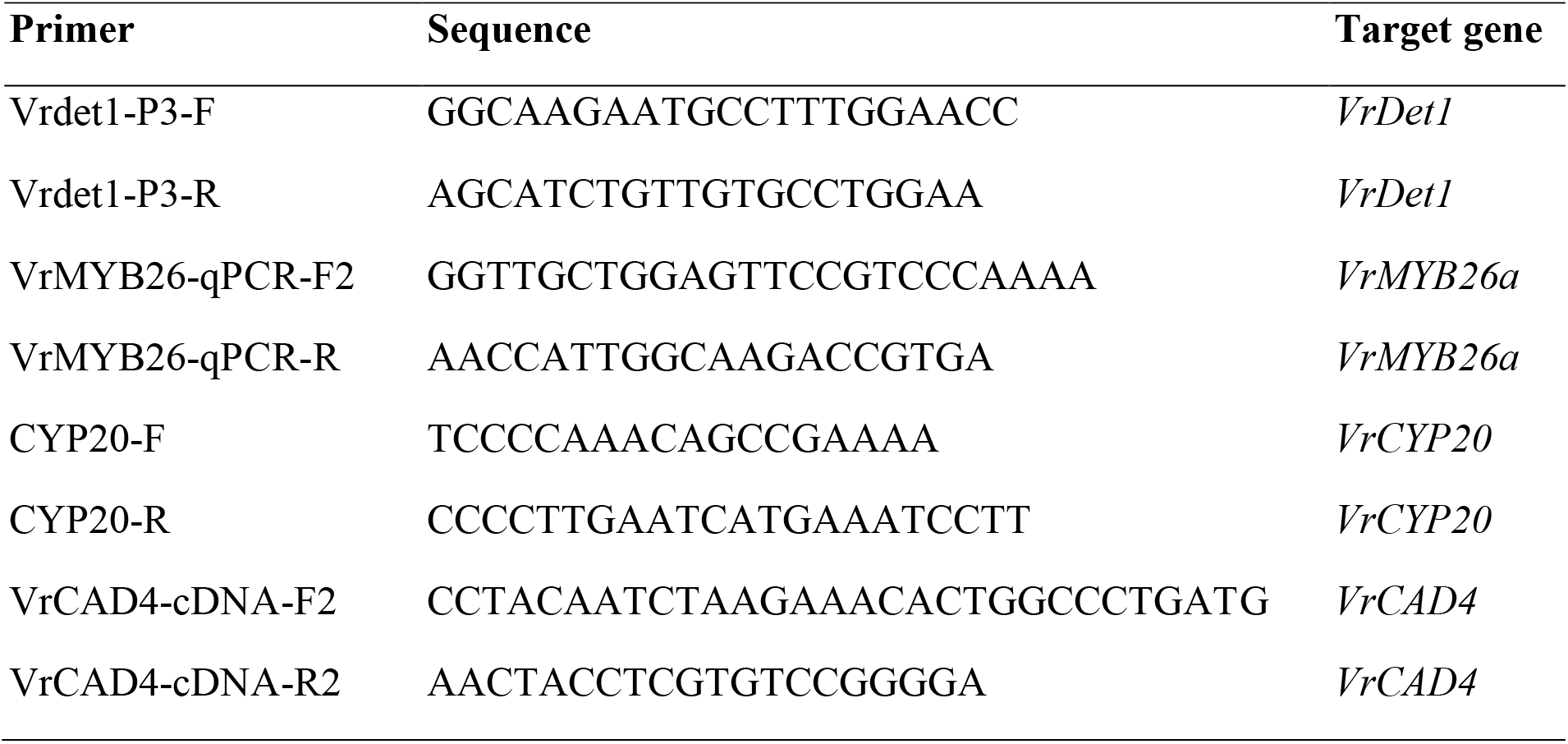
List of primers used in this study.

